# Functional specification of CCK+ interneurons by alternative isoforms of Kv4.3 auxiliary subunits

**DOI:** 10.1101/683755

**Authors:** Viktor János Oláh, David Lukacsovich, Jochen Winterer, Andrea Lőrincz, Zoltan Nusser, Csaba Földy, János Szabadics

## Abstract

CCK-expressing interneurons (CCK+INs) are crucial for controlling hippocampal activity. We found two firing phenotypes of CCK+INs in rat CA3 area; either possessing a previously undetected membrane potential-dependent firing or regular firing phenotype, due to different low-voltage-activated potassium currents. These different excitability properties destine the two types for distinct functions, because the former is essentially silenced during realistic 8-15 Hz oscillations. The general excitability, morphology and gene-profiles of the two types were surprisingly similar. Even the expression of Kv4.3 channels were comparable, despite evidences showing that Kv4.3-mediated currents underlie the distinct firing properties. Instead, the firing phenotypes were correlated with the presence of distinct isoforms of Kv4 auxiliary subunits (KChIP1 vs. KChIP4e and DPP6S). Our results reveal the underlying mechanisms of two previously unknown types of CCK+INs and demonstrate that alternative splicing of few genes, which may be viewed as a minor event in the cells’ whole transcriptome, can underlie distinct cell-type identity.

## Introduction

The biophysical and morphological properties of complementary GABAergic cell classes are specifically tuned for regulating the activity of the much more populous principal cells in broad temporal (from second to sub-millisecond range; Hu et al., 2014, Overstreet-Wadiche and McBain, 2015) and spatial domains (from axons to distal dendrites; Freund and Buzsaki, 1996). Thus, specialized features of GABAergic neurons help the hippocampus to comply with the vast computational demand related to various behaviors (Klausberger and Somogyi, 2008). However, the currently known degree of functional diversity of GABAergic cells cannot match the vast amount of hippocampal behavioral tasks. Therefore, a more complete understanding of the diversity of GABAergic neurons is one of the major goals of current research. Indeed, emerging evidences from new technologies that map the complete transcriptome of individual cells suggest that the number of GABAergic types may be higher than currently recognized (Földy et al., 2016, Fuzik et al., 2016, Harris et al., 2018, Que et al., 2019, Tasic, 2018, Zeisel et al., 2015). These new data sets accumulate a deep understanding of cellular diversity at the level of genes, which requires the subsequent determination of the relation between genes and higher level complexities, such as cellular excitability or morphology, which ultimately defines the distinct physiological functions of individual cells. Thus, multidimensional approaches are needed for establishing links between gene level diversity and the functional contribution of identified cell types. However, unanswered questions prevent us from establishing links between these components. Is the diversity of different gene profiles sufficient for different function, or the multitude of changes converge into similar phenotypes (Marder and Goaillard, 2006)? How many genes are needed for different cellular functions? To better understand the boundaries of cell classes, here we employed various approaches to describe a new level of functional diversity among CCK-expressing hippocampal interneurons (CCK+IN) in rat area CA3 and link this to specific single cell level differences in gene profiles, expression of proteins and ionic currents.

Several properties distinguish CCK+INs from other major GABAergic cell classes. Unlike in other GABAergic cells, the axons of CCK+INs are highly enriched with CB1 receptors, which mediate activity-dependent regulation of individual synapses (Freund and Katona, 2007). As a result of this delicate feedback control, CCK+INs are ideally suited for dynamic inhibition of a subset of principal cells based on the context of ongoing activity. The *in vitro* firing of CCK+INs is believed to be homogeneous, regular and distinct from other classes (Cea-del Rio et al., 2011, Glickfeld and Scanziani, 2006, Szabadics and Soltesz, 2009, Szabo et al., 2014). The conventional view of CCK+IN firing includes intermediate AP widths (between that of pyramidal cells and classical fast-spiking interneurons) and clear spike frequency accommodation. These excitability properties are crucial for the generation of distinct firing of CCK+INs that is observed during exploration associated with theta and gamma oscillations (Klausberger et al., 2005, Lasztoczi et al., 2011) in response to specific synaptic inputs (Glickfeld and Scanziani, 2006, Matyas et al., 2004). However, the CCK+IN class is diverse. Based on their axonal morphology basket-, mossy fiber-associated, Schaffer collateral-associated and perforant path-associated types (Cope et al., 2002, Vida and Frotscher, 2000, Vida et al., 1998) can be distinguished. These distinct morphological types selectively control excitations from various sources (hence their names). Furthermore, there are molecules that are complimentarily expressed by subsets of CCK+INs, such as VGluT3 and VIP (Somogyi et al., 2004) and single-cell RNA-sequencing also uncovered several additional molecular varieties (Fuzik et al., 2016). Thus, due to their distinction from other major classes and large interclass variability, CCK+INs are ideal for addressing general questions concerning the relation of genes to cellular identity and boundaries between GABAergic cell classes.

In this study, we investigated hippocampal CCK+INs from a broader perspective that could reveal previously undetected excitability parameters suitable for specific physiological functions. A hint that such diverse physiological parameters exist came from previous *in vivo* recordings that showed that individual CCK+INs are differentially active during various oscillatory states (Klausberger et al., 2005, Lasztoczi et al., 2011), where many of them appeared to prefer either lower or higher frequency ranges. We found two types of hippocampal CCK+INs in rats based on their different excitability, with potentially different contributions to network events, particularly in the range of theta oscillations. Detailed realistic simulations showed that switching only the properties of Kv4.3-mediated currents can sufficiently convert one functional cell type to the other. Combined analyses of the complete mRNA content and protein expression of single cells revealed that the pronounced functional distinction between the here-described two CCK+IN type is defined by differential isoform usage of three auxiliary subunits of the Kv4.3 channels.

## Results

### Half of the CA3 CCK+INs show state-dependent firing

To explore potential differences in the excitability of individual CCK+INs in the CA3 region of the rat hippocampus, first we characterized their firing properties in two different conditions. Specifically, we recorded their spiking in response to current steps from two, physiologically plausible membrane potential ranges of *post hoc* identified CCK+INs. We focused mostly on CA3 region because CA3 CCK+INs are the most diverse within the hippocampus. When CCK+INs (n = 557 cells) were stimulated from slightly depolarized membrane potentials (MP, range: -60 – -65 mV) relative to rest (−64.7 ± 0.4 mV), action potential (AP) firing always showed spike-frequency accommodation, which is one of the most characteristic features of this cell class (Cea-del Rio et al., 2011, Glickfeld and Scanziani, 2006, Szabadics and Soltesz, 2009, Szabo et al., 2014). However, we noticed that numerous CCK+INs (n = 290 cells) showed MP-dependent firing: their initial spiking was strongly inhibited and its onset was delayed when it was evoked from hyperpolarized MPs (between -75 to -85 mV, Figure 1A-B). On average, these cells started firing after a 252 ± 15 ms silent period from hyperpolarized MP (measured from the start of the current injection). We categorized these cells as Transient Outward Rectifying cells or TOR cells (a term that was used to describe cells in other brain regions: Stern and Armstrong, 1996). The rest of CCK+INs (n = 267 cells) were characterized as regular spiking or RS cells as they fired regularly irrespective of their MP and they started firing with a short delay (33 ± 2 ms) when stimulated from hyperpolarized MP. At depolarized MP (−55 to -65 mV), the first APs of both TOR and RS cells occurred with short delays (48 ± 3 ms and 26 ± 1 ms, respectively, *Student t-test, p = 0.09, t(160) = -1.706*).

**Figure 1.**
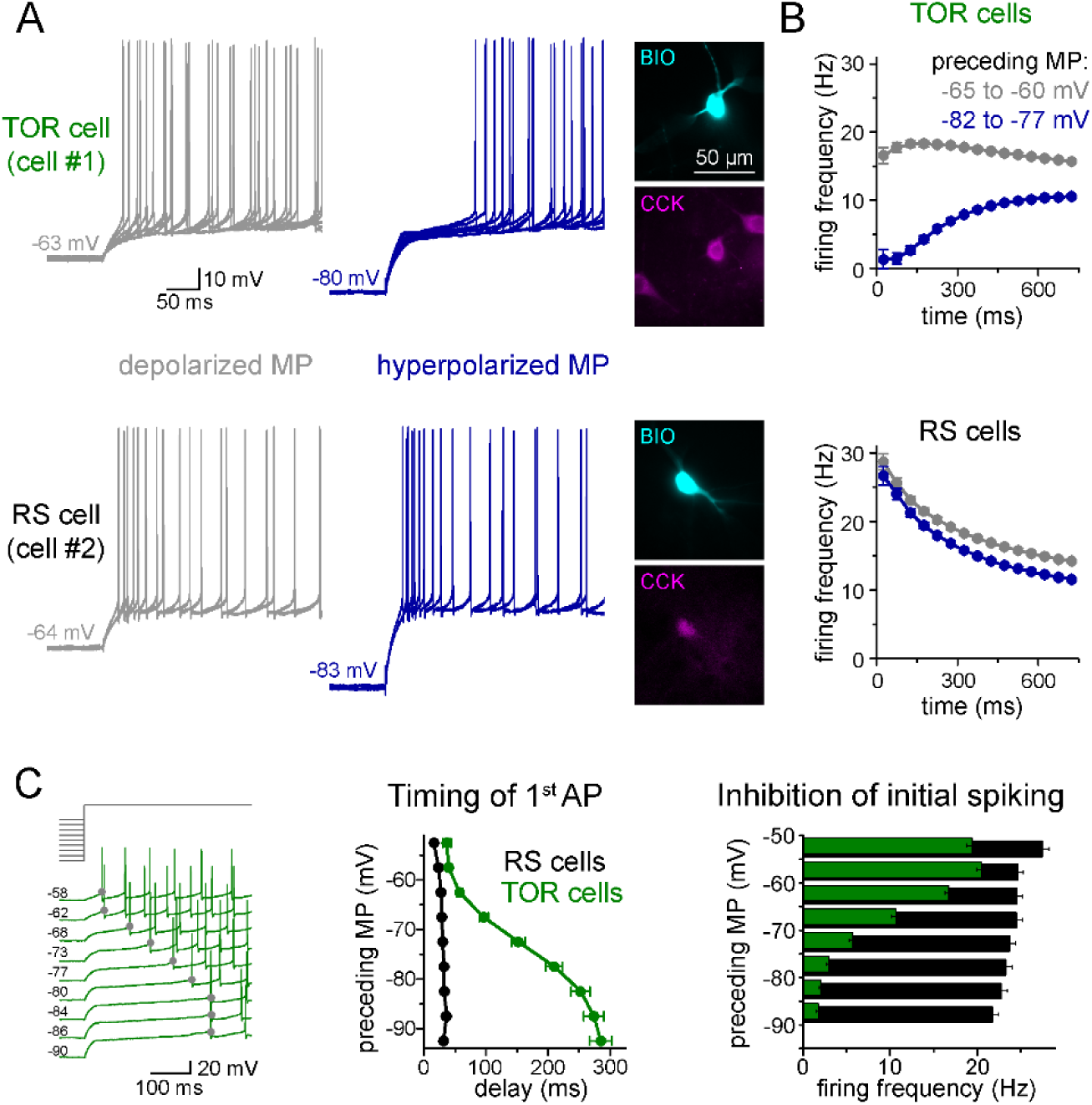
Two distinct firing patterns within CA3 CCK+ cells. **A**. Firing properties of two representative CCK+INs in the CA3 hippocampal region. Firing was elicited with square pulse current injection of identical amplitude, but from depolarized (grey traces), or hyperpolarized MPs (blue traces). Several trials are superimposed to show the stability of the timing of the first action potential. Insets show the immunolabelling of the biocytin filled (BIO) recorded cells for CCK. **B**. Average time course of AP occurrence in TOR and RS cells from two MP ranges (n = 120 and 113 representative cells, respectively). **C**. Timing of the first AP and probability of APs during the first 150 ms of the square pulse stimulus shows steep MP-dependence in TOR cells, whereas the initial spikes are stable in the RS cells. The amplitude of stimulating current steps was standardized for each cell and only the preceding holding current (3 seconds) was varied in individual trials. Traces show a representative recording from a TOR cell. The average data derived from 85 TOR and 81 RS cells.

Next, we applied a protocol allowing the detailed quantification of the MP-dependence of firing in individual cells (n = 81 RS and 85 TOR cells). Specifically, firing was evoked by a current step that was calibrated for each cell to elicit similar average firing (10-20 Hz) from slightly depolarized MP. The holding current preceding the standard step was systematically varied to reach a wide range of steady-state MPs (3 seconds, in the range of -50 and -90 mV, Figure 1C). In TOR cells, the number of APs within the first 150 ms and the timing of the first AP showed steep voltage dependence (V_1/2_ value of the Boltzmann fits were -67.4 mV and -73 mV, respectively, R^2^= 0.995 and 0.999, Figure 1C). In contrast, the delay and number of spikes did not show membrane potential-dependence in RS cells.

Unlike the initial spiking, the general firing and membrane properties of TOR and RS cells were similar, including input resistance (143 ± 7 MΩ vs. 139 ± 7 MΩ, n = 112 TOR and n = 122 RS cells, respectively), AP threshold (−37.4 ± 0.4 mV vs. -37.6 ± 0.4 mV), AP half-width (0.49 ± 0.01 ms vs. 0.51 ± 0.01 ms), *d*V/*d*t maximum (484 ± 13 mV/ms vs. 486 ± 14 mV/ms) and AHP amplitude (−15.5 ± 0.3 mV vs. -14.4 ± 0.3 mV). Furthermore, the frequency of spontaneous synaptic events (including both IPSCs and EPSCs; 2.47 ± 0.57 Hz and 3.26 ± 0.86 Hz, n = 9 TOR and 8 RS cells) and amplitude of these events (−51.6 ± 3.3 pA and -51.5 ± 2.5 pA) were also similar. TOR cells were present in the CA1 region as well. However, here their prevalence was lower compared to CA3 (2 TOR out of 13 CA1 CCK+IN). Both TOR (n = 15 cells) and RS (n = 12 cells) CCK+INs were also detected in the CA3 region of adult rats (older than 70 days). In summary, the firing of TOR cells shows a remarkable sensitivity to a physiologically plausible 20 mV shift in the MP, despite having no other distinctive passive electrical or spiking properties compared to RS cells.

### TOR and RS firing types do not correlate with previously known subtypes of CCK+ cells

The CCK+IN class has been previously divided into several subtypes based on various functionally relevant features. Therefore, next we investigated whether the two MP-dependent firing phenotype can be linked to previously known subtypes of CCK+INs. Three subtypes have been previously identified within CA3 CCK+INs based on the target zones of their axons. Basket cells (BCs) innervate the soma and proximal dendrites (Hendry and Jones, 1985). Mossy fiber-associated (MFA) cells are specific to the CA3 region where their axons colocalize with the axons of dentate gyrus granule cells in stratum lucidum and hilus (Vida and Frotscher, 2000). The third morphological type is named after the position of their axons as Schaffer collateral associated cells (SCA)(Cope et al., 2002). Altogether, 172 of the recorded CCK+INs were unequivocally identified either as BCs (n = 31), MFAs (n = 96) or SCAs (n = 45; Figure 2A). Both TOR and RS firing types occurred similarly among these morphological subtypes (Figure 2B) indicating that the two firing phenotypes cannot be assigned to these morphology-based subtypes of CCK+INs.

**Figure 2.**
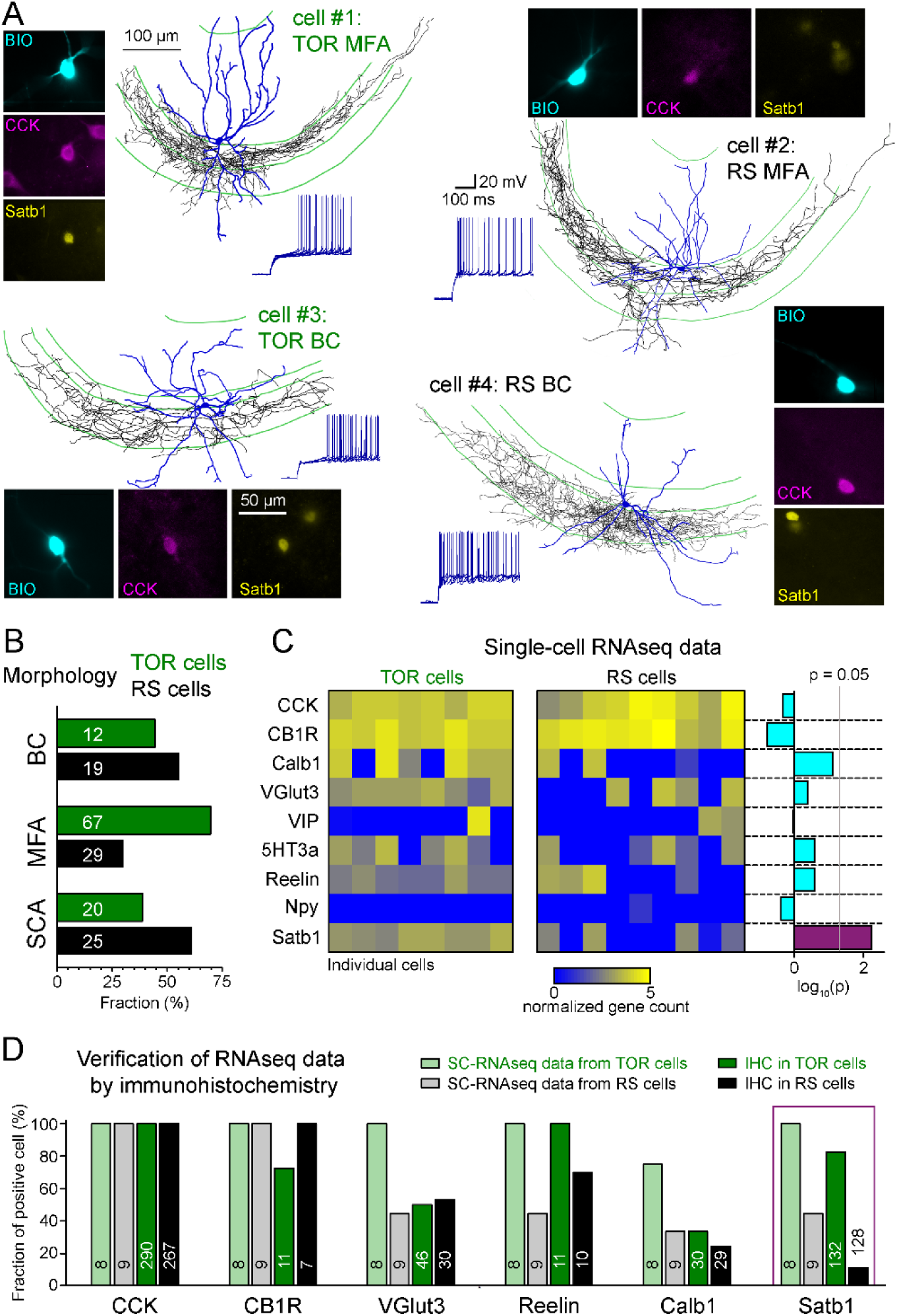
Different excitability does not correlate with previously known diversity of CCK+ cells. **A**. Axonal (black) and dendritic (blue) reconstructions of four CCK+INs in the CA3 region representing two major morphological types, the mossy-fiber associated cells and basket cells. The firing of cell#1 and #2 are shown in detail in Figure 1A. The firing from hyperpolarized MP is shown for each cell. Immunolabellings for CCK and the nuclear protein Satb1 are shown next to each recorded cell. **B**. Prevalence of TOR and RS cells within three major morphological types of CA3 CCK+. Numbers of identified cells are indicated on each bar (BC: CCK+ basket cells, MFA: mossy-fiber associated cells, SCA: Schaffer-collateral associated cells). **C**. Single cell RNAseq characterization of recorded TOR (left, n = 8) and RS (right, n = 9) cells for known GABAergic cell markers. Each column corresponds to single identified CCK+INs. Only Satb1 mRNA content was significantly different between TOR and RS cells (*p = 0.00346, Mann-Whitney Test*). **D**. Comparison of immunhistochemical and single cell RNAseq data. Bar plots show the fraction of the recorded cells with detectable RNA content (threshold: 0.2) for the selected markers and immunopositivity for the same proteins (for examples of the immunolabelling see Figure 2 – figure supplement 1). The number of tested cells are shown on each bar.

In addition to the variable axonal morphology, CCK+INs are known to heterogeneously express several molecules (Somogyi et al., 2004). To compare their molecular content first we performed single cell RNA sequencing (SC-RNAseq) (Földy et al., 2016) of individually recorded TOR cells (n = 8) and RS cells (n = 9). Only those cells were analyzed in detail, which had detectable mRNA for both CCK and cannabinoid-receptor type 1 (CB1R or Cnr1; Figure 2C). In accordance with previous observations (Földy et al., 2016), the included cells contained the transcripts of at least 3000 different genes. Most subtype-specific marker genes (calbindin/Calb1, vesicular glutamate transporter VGlut3/slc17a8, vasoactive intestinal protein/VIP, serotonin receptor subtype 3a/5HT3a, Reelin, neuropeptide-Y/Npy) were present at comparable levels in TOR and RS cells (Figure 2C), except the activity-dependent transcription regulator, Satb1 (Close et al., 2012). All tested TOR cells had high levels of Satb1 mRNA, whereas it was present only in 4 of the 9 tested RS cells (*Mann-Whitney test, p = 0.0058, z = 2.76, U = 65*). Furthermore, to compare the general gene profiles, we performed hierarchical cluster analysis based on all genes that were detected in at least 3 of the 17 tested cells (10095 out of 30662 tested genes, data not shown). This analysis did not categorize these cells into two distinct groups that would correspond to their firing phenotype, arguing for their similar cell class identity (Fuzik et al., 2016, Harris et al., 2018).

**Figure 2 – figure supplement 1.**
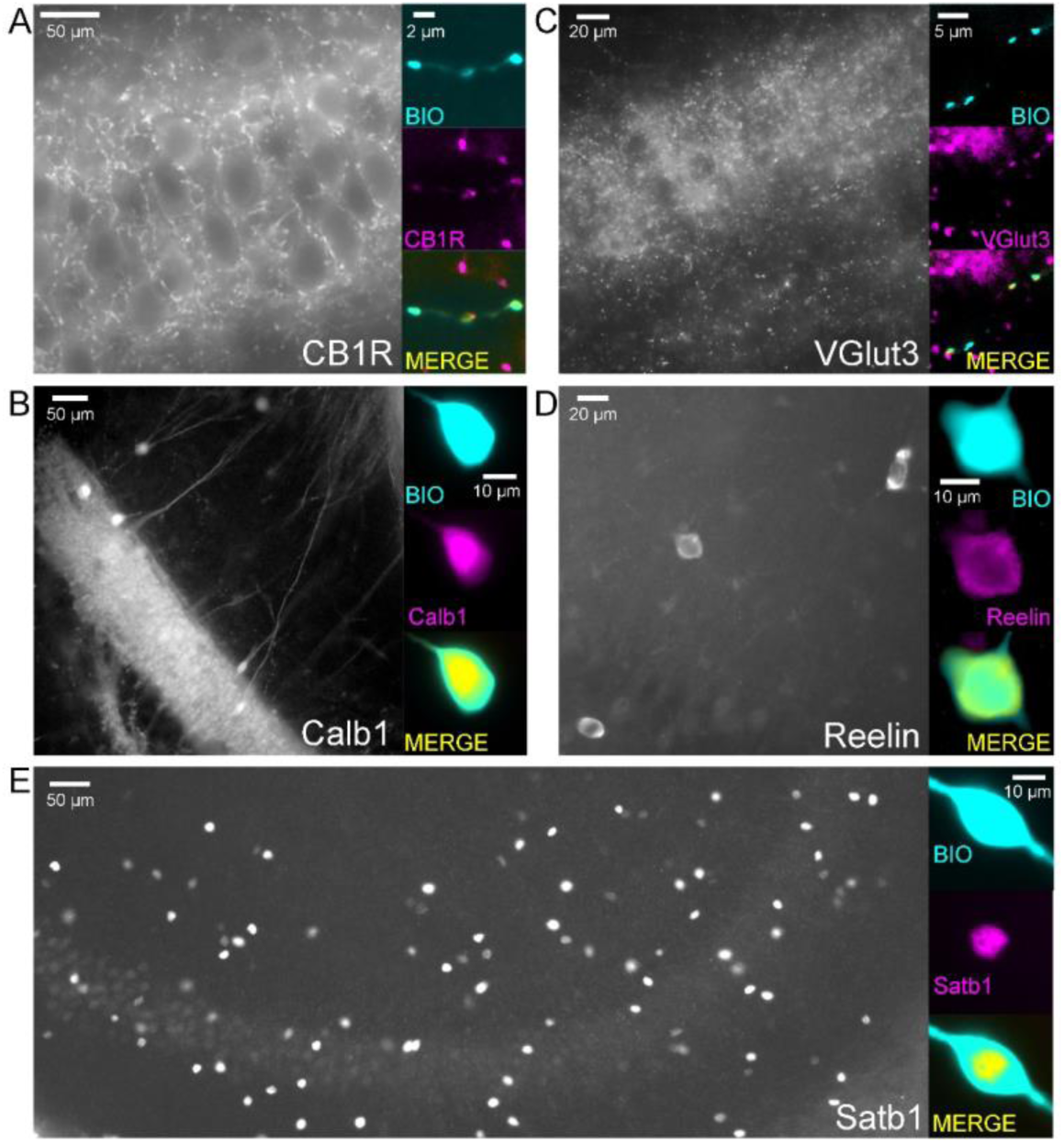
Example cells that were immunopositive for CB1R (**A**), Calb1 (**B**), VGluT3 (**C**), Reelin (**D**) and Satb1 (**E**) and the low magnification overview of the labeling pattern of these molecules within the CA3 region.

To confirm these findings at the level of protein expression, next we performed immunohistochemistry on identified CCK+ TOR and RS cells in separate experiments (Figure 2D and Figure 2 - figure supplement 1). As expected, CB1R protein was detected in the axon of almost all tested cells (15 out of 18 tested cells). Furthermore, VGluT3 (23 out of 46 TOR cells, 16 out of 30 RS cells), Reelin (all TOR, n = 11 and 7 out 10 RS was positive) and Calb1 (10 out 30 TOR and 7 out of 29 RS) proteins occurred similarly in RS and TOR types. However, the majority of the tested TOR cells were positive for Satb1 protein (109 out of 132, 82.6%) but only few RS cells were positive (14 out of 128 tested, 10.9%). Satb1 protein was also prevalent in non-recorded CCK+ cells within the CA3 region of the acute slices (126 Satb1+/CCK+ out of 425 CCK+), suggesting that the presence of this activity-dependent marker is not due to the recording. The somata of Satb1+/CCK+ cells were found throughout CA3 strata oriens, pyramidale, lucidum and radiatum and their proportion to all CCK+ cells were variable (34.1%, 21.4%, 33.6%, and 27.2%, respectively). In agreement with the lower occurrence of TOR cells in the CA1, non-recorded CCK+ cells in this region were less likely to be Satb1 positive (21.6% of 231 CCK+ cells). In the hilar region of the dentate gyrus the proportion of Satb1+/CCK+ cells was also low (4.8% of 123 CCK+ cells). We have also detected Satb1 and CCK overlap in tissue samples derived from transcardially perfused animals (20.2%, 97 Satb1+/CCK+ out of 480 CCK+ cells in the CA3 area). Thus, the mRNA and immunohistochemical data suggest that the Satb1 content of hippocampal CCK+INs is predictive for their TOR or RS identity and these firing phenotypes are not related to previously known subtypes of the CCK+IN class.

### Differences in low-voltage-activated potassium currents (ISA) underlie the heterogeneity of CCK+IN firing

Next, we investigated the conductances that are responsible for the difference of the two firing types in CCK+INs. We focused on near-threshold potassium currents, which can effectively regulate firing responses in similar experimental conditions in various brain regions (Lammel et al., 2008, Margrie et al., 2001, Neuhoff et al., 2002, Stern and Armstrong, 1996). First, we characterized each cell as TOR or RS type under normal recording conditions. Then, we blocked sodium (3 µM TTX) and hyperpolarization-activated cation currents (10 µM ZD7288) and recorded near-threshold potassium currents by applying 300 ms voltage command steps between -100 and -25 mV from -120 mV conditioning potentials (300 ms). All cells included in the analyses were verified to be CCK+ by *post hoc* immunolabelling. TOR cells had a substantial amount of potassium currents at firing threshold (Figure 3A, 683 ± 109 pA at -40 mV). In contrast, potassium currents in RS cells activated at more positive voltages and were much smaller at threshold (148 ± 28 pA, *p = 0.0006, t(26)=3.885, n = 17 and 11, Student’s t-test*). Because in these recording conditions the majority of the membrane conductances remained intact, voltage clamping could not be properly performed at more depolarized MPs, which precluded the determination of the exact half-activation voltage values. The robust low-voltage-activated potassium currents (I_SA_), which are activated tens of millivolts below AP threshold, can underlie the strong inhibition of AP generation in TOR cells whereas I_SA_ in RS cells is much smaller at near-threshold voltage range.

**Figure 3.**
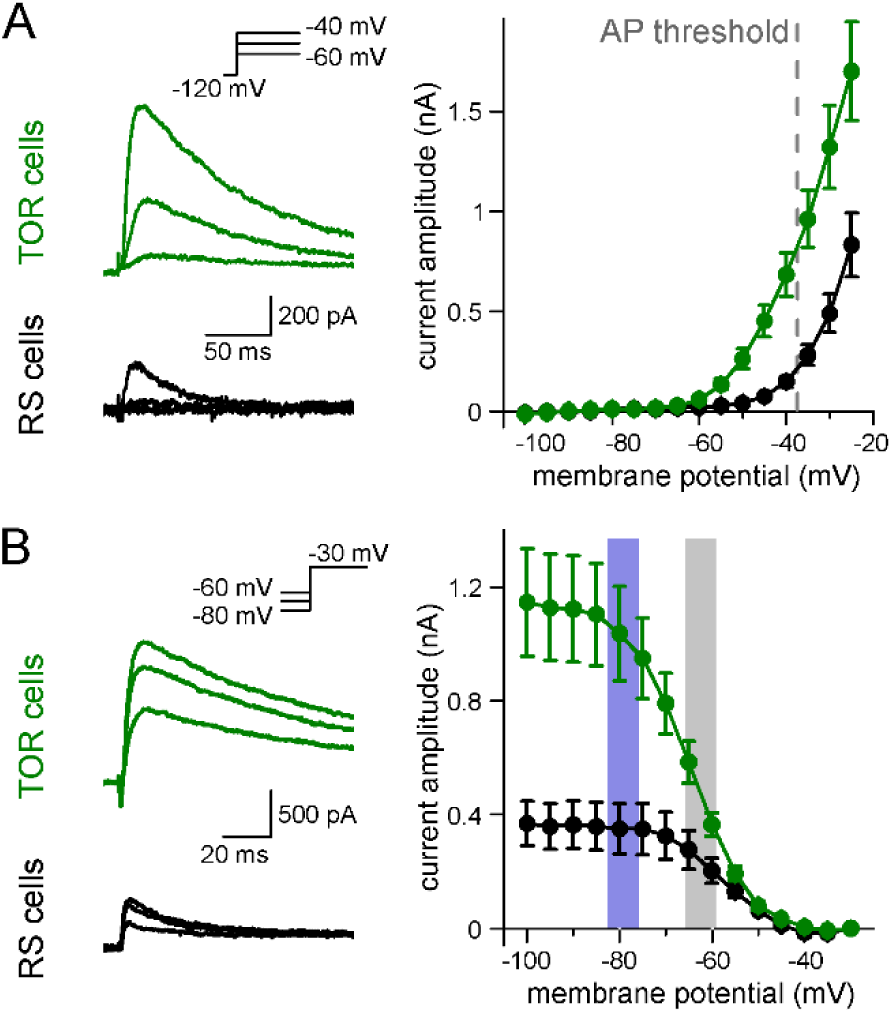
Differences in I_SA_ currents underlie the heterogeneity of CCK+IN firing. **A**. Representative traces of low voltage activated potassium currents from TOR and RS cells (in the presence of 3 µM TTX and 10 µM ZD7288). Right, voltage dependence of activation of I_SA_ in TOR (n = 17) and RS (n = 11) cells. The grey dotted line indicates AP threshold (−37.52 ± 0.3 mV) measured before TTX application. Notice the large amount of outward current in TOR cells at subthreshold MPs. **B**. Representative traces of I_SA_ activated at -30 mV from different holding potentials. Right, voltage dependence of inactivation of I_SA_ in the two cell types (n = 14 and 10 for TOR and RS cells respectively). Blue and grey shaded areas indicate the voltage ranges from which state-dependent firing was tested (see Figure 1).

Next, we investigated the availability of I_SA_ at different MPs. For these measurements, potassium currents were evoked by a voltage step to -30 mV following various pre-pulse potentials between -100 to -35 mV. The V_1/2_ of the average inactivation curve was -64.5 ± 0.2 mV (*Boltzmann fit, R*^*2*^ *= 0.999*, mean of V_1/2_ from individual cells: -63.8 ± 0.9 mV, *n = 13 cells*, Figure 3B) in TOR cells. Importantly, the majority of I_SA_ was available at slightly hyperpolarized MPs (91.3 ± 1.6% at -80 mV), where the inhibition of firing was clearly observable during the characterization of TOR cells (see Figure 1 data). But at -60 mV, where the inhibition of spiking was not prominent, the majority of outward currents in TOR cells were inactivated; only 35.7 ± 3.4% of the current was available. Thus, the MP-dependence of the steady-state inactivation of I_SA_ can explain the TOR firing phenotype. Similar to the activation, the inactivation of I_SA_ in RS cells was shifted toward positive voltage ranges (V_1/2_: -57.4 ± 0.3 mV, n = 8 cells, mean of individual data: -55.6 ± 2 mV, comparison with TOR cells: *p = 0.0006, t(19)=4.12, Student’s t-test*) and a larger portion of this smaller current was available at -60 mV (52.7 ± 6.3%, Figure 3B). Thus, hyperpolarization of the RS cells cannot add a substantial amount of AP-firing disabling inhibitory conductance.

Interestingly, the inactivation time constant of I_SA_ was faster in RS cells compared to TOR cells (18.8 ± 1.7 ms vs. 71.2 ± 9.1 ms, measured at -25 mV, *p = 0.0007, t(19)=4.025, Student’s t-test*, n = 7 and 14). Furthermore, the recovery from inactivation was faster in RS cells, but it showed steep voltage dependence in TOR cells (TOR cells: time constants of the recovery were 58.2 ± 2.9 ms, 46.2 ± 2.9 ms, 29.8 ± 2.5 ms and 4.4 ± 0.1 ms at -65, -75, -85 and -120 mV, respectively; whereas in RS cells: 11.8 ± 4.4 ms, 8.0 ± 2.2 ms, 7.1 ± 1.6 ms and 3.3 ± 0.4 ms, respectively). Due to these properties, I_SA_ currents in RS cells resulted in only a small inhibitory charge transfer around the AP threshold, whose availability remains similarly limited within the relevant -80 and -60 mV MP range. Thus, the different properties of I_SA_ currents, particularly the left-shifted inactivation and activation curves, can explain the differences in TOR and RS firing phenotypes.

### Realistic models of TOR and RS firing

In the previous experiments, we injected simple steady-state currents to CCK+INs, which already revealed remarkably different excitability properties. However, excitation of neurons *in vivo* is more dynamic due to the fast rise and decay of PSPs that are clustered into packages according to the ongoing oscillatory state of the network, resulting in a constantly fluctuating membrane potential. The different temporal properties of I_SA_ in TOR and RS suggest that these cells can follow membrane potential fluctuations differently. Therefore, to predict the frequency ranges of oscillations that are optimal for the distinct CCK+IN types with distinct excitability, we performed computer simulations of realistic dynamic behavior of RS and TOR phenotypes.

First, we equipped five reconstructed CCK+INs (3 TOR and 2 RS types) with known voltage-dependent conductances and passive properties of hippocampal CCK+INs (Bezaire et al., 2016) and reproduced the general firing properties (Figure 4A). To add the I_SA_ currents to the models, we recorded the properties of local I_SA_ current by pulling outside-out patches from the soma and dendrites of RS and TOR cells (Figure 4 – figure supplement 1A-B). This additional experiment was necessary because the subcellular distribution of many ion channels are inhomogeneous (Nusser, 2009), which also influences the somatically measured ensemble currents. I_SA_ was isolated by subtraction of currents measured at 0 mV with -80 and -50 mV pre-pulses. We did not observe a significant gradient along the dendritic axis of the two cell types, albeit there was a tendency for larger somatic current densities compared to dendrites. The average kinetics of patch currents matched those of whole-cell currents: slower inactivation in TOR cells/patches. However, we detected a large variability of I_SA_ between individual patches from TOR cell (Figure 4 – figure supplement 1C), even when multiple patches were pulled from the same cell. Patches from RS cells were more homogeneous (variance of current decays, TOR: 2888 ms^2^, RS: 373 ms^2^). Surprisingly, in many TOR cell patches the current kinetics resembled the patch- and whole-cell currents of the RS cells. While in other TOR cell patches much slower currents were also detected. This variability might be due to the clustered occurrence of voltage-gated potassium channels in GABAergic cells (e.g. see Kollo et al., 2006). We implemented this variability by equipping TOR models with two types of I_SA_ currents (for details see below).

**Figure 4.**
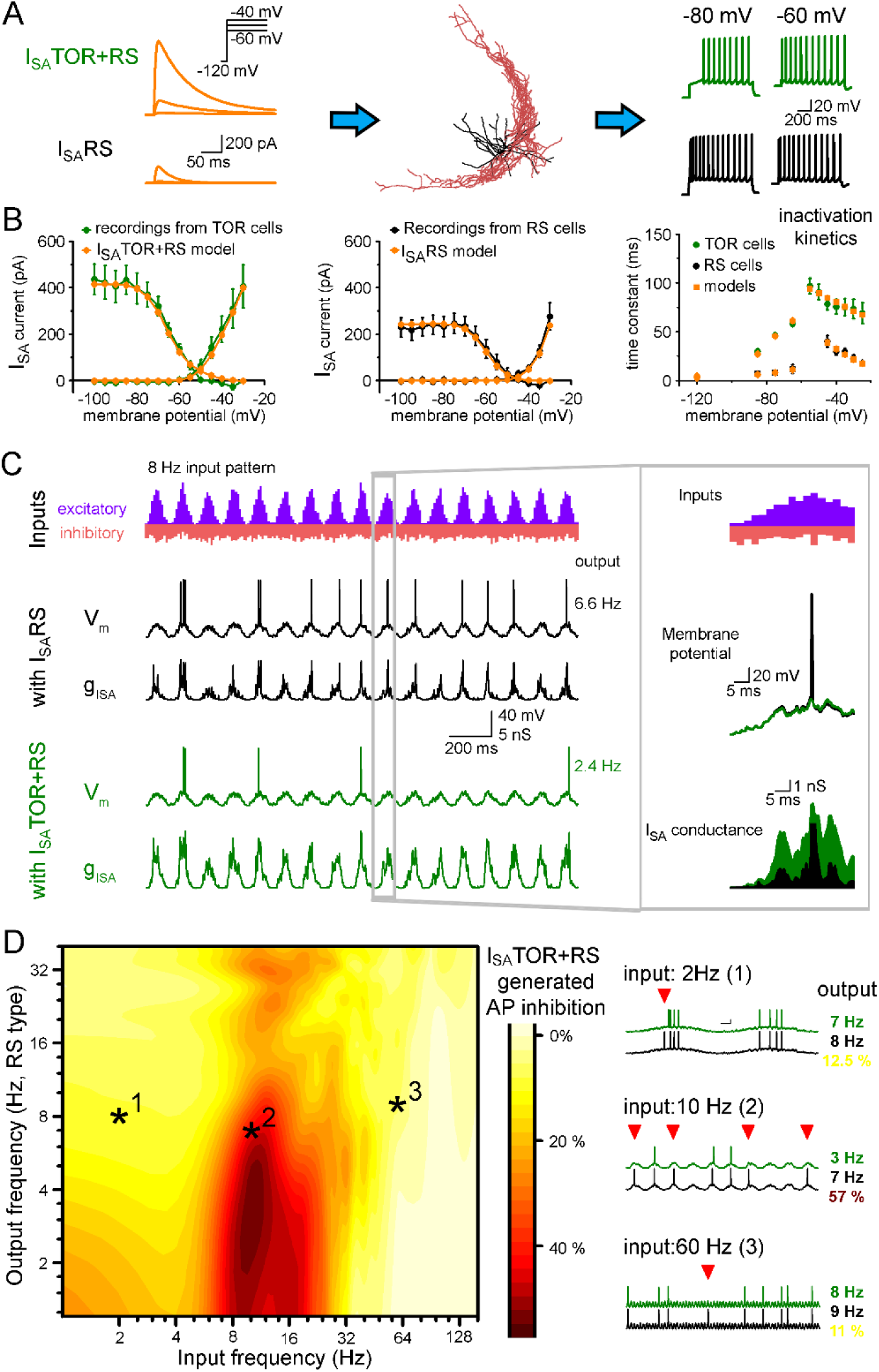
Different I_SA_ currents in TOR and RS cells tune them for different network states. **A**. I_SA_TOR+RS and I_SA_RS were added to five reconstructed CCK+INs, which possess every known voltage-dependent conductance (Bezaire et al., 2016). Changing I_SA_RS to I_SA_TOR+RS in the same cells transformed the firing from RS to TOR phenotype. These simulated cells were equipped with synaptic conductance to simulate input drives in various network states. **B**. Voltage clamp simulations with complete morphology and realistic conductances reproduced whole-cell I_SA_ currents of TOR and RS cells, including voltage-dependence, the total current measured at the soma (left and middle graphs), and inactivation and the kinetics of inactivation and recovery from inactivation (right graph, where recovery was measured at -120, -85, -75 and -65 mV). Note that the total amount of I_SA_ in RS cells was larger than in TOR cells (Figure 4 – figure supplement 1A-C) and the larger current at low voltages are the consequence of the left-shifted activation curve. **C**. Representative traces showing the number and temporal distribution of 8 Hz-modulated synaptic inputs a simulated CCK+INs and its MP and I_SA_ conductance in two conditions with either I_SA_RS or I_SA_TOR+RS. **D**. Average effects of exchanging I_SA_RS to I_SA_TOR+RS on the output of five CCK+INs during various input frequency ranges (x-axis) and baseline output activity (i.e. with I_SA_RS conductance, y-axis). Yellow color shows no change in firing when I_SA_TOR+RS replaced I_SA_RS, whereas red color indicates robust reduction in AP output. Representative traces on the right depict three examples with different input frequencies. Red triangles highlight inhibited spikes.

**Figure 4 – figure supplement 1.**
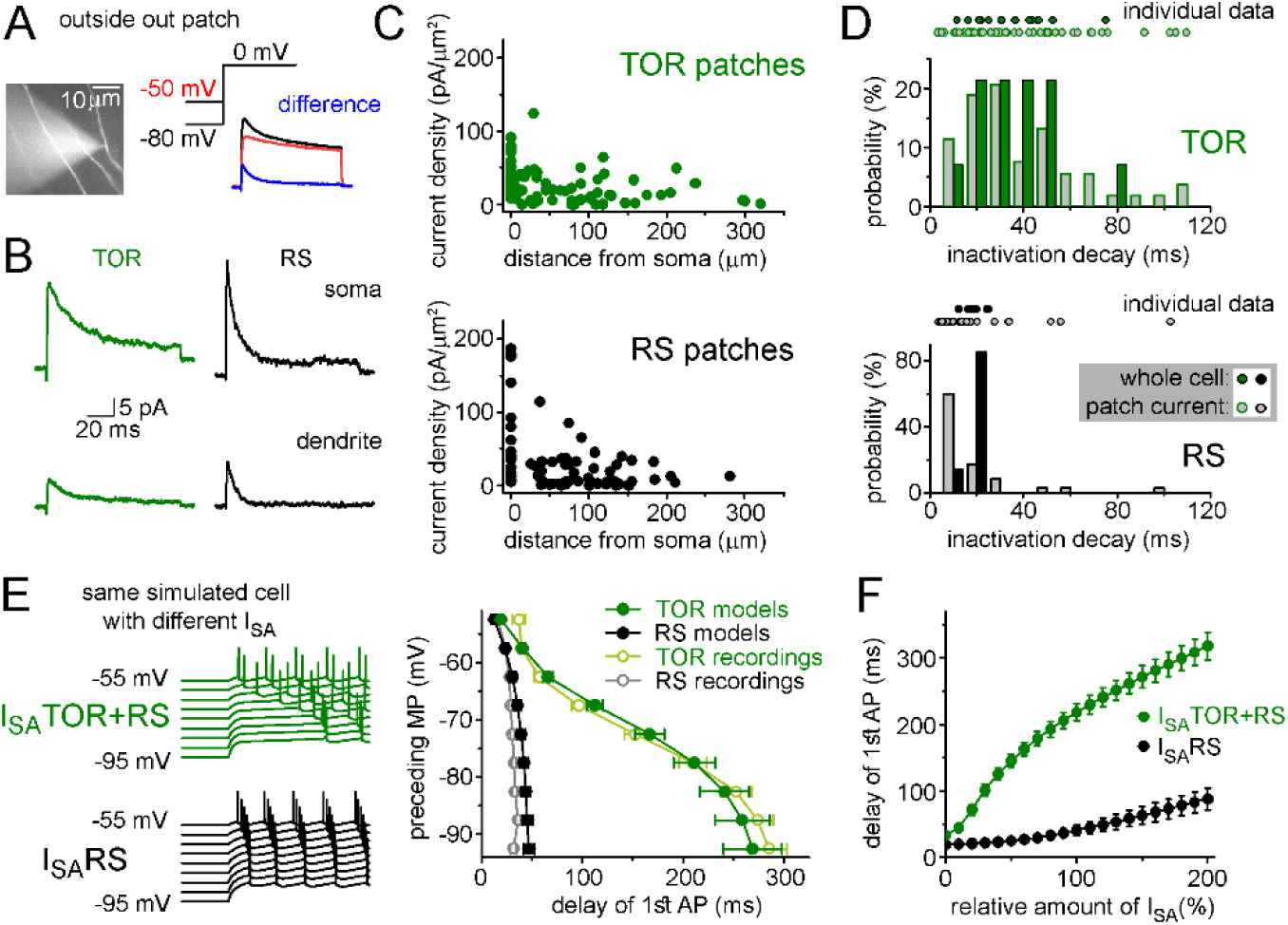
Realistic CCK+INs in-silico. **A**. Representative outside-out patch recording of dendritic I_SA_ current from a RS cell, which was somatically loaded with Alexa594 dye to visualize dendrites in epifluorescent illumination and to record firing pattern. I_SA_ current (blue) was evoked at 0 mV and isolated by subtracting currents evoked with -80 mV prepulse voltage from -50 mV prepulse. **B**. Average I_SA_ currents from somata and dendrites of RS and TOR cells (n = 22 somatic TOR-, n = 17 somatic RS-, n = 56 dendritic TOR and n = 53 dendritic RS patches). **C**. Current densities are plotted against dendritic distance. Current densities were calculated by dividing the peak of I_SA_ current with the membrane surface of the patch, which was calculated from the capacitance difference measured with the patch membrane and after pushing the pipette in an insulator gel (Sylgard). **D**. Comparison of inactivation decay time constants of I_SA_ measured in individual whole-cell and outside-out recordings (TOR patch: 74.5 ± 7.4 ms, variance: 2888, n = 53; TOR whole-cell, 71.2 ± 9.1 ms, variance: 1147, n = 14, RS patch: 17.2 ± 3.3 ms, variance: 373, n = 35). Symbols above the graphs show individual measurements and their distributions are represented by bar graphs. Notice the wider distribution of patch data compared to whole-cell data from TOR cells. **E**. Left, representative current-clamp simulations from a single model cell with either I_SA_TOR+RS or I_SA_RS currents. Notice the characteristic MP-dependence of spike onset with I_SA_TOR+RS. The graph on the right summarizes the MP-dependence of timing of the first APs from five model cells with either I_SA_TOR+RS or I_SA_RS. The results from recordings using the same protocols are shown for comparison (from Figure 1C, open symbols). **F**. Conductance density has little effect on the presence or absence and the extent of the TOR phenomenon suggesting that the kinetic and voltage-dependent properties of the two sets of I_SA_ currents play a more important role in this phenomenon than the amounts of inhibitory conductance (measured from -80 mV preceding MP).

These outside-out patch recordings also allowed us to directly compare the density of I_SA_ in TOR and RS cells because the majority of channels are expected to be open at larger voltage steps (to 0 mV) and the current amplitude is determined by the driving force and the conductance. Interestingly, the density of somatic I_SA_ in RS cells were significantly larger compared to that of the TOR cells (70.5 ± 15.2 pA/µm^2^ vs. 42.2 ± 5.2 pA/µm^2^, n = 26 and 17, *p = 0.047, t(41)=-2.048*). This result does not contradict with the above data that the total I_SA_ current is larger in TOR cells near the AP threshold because in TOR cells the activation of I_SA_ is left-shifted, therefore, a larger fraction of channels is open at lower voltages. In dendritic patches (25-320 µm) the density of I_SA_ was similar in TOR and RS cells (19.6 ± 3.3 pA/µm^2^ vs. 19.5 ± 3.0 pA/µm^2^, n = 53 and 47).

Based on the results of the above recordings, we used two sets of I_SA_ conductances and tuned their densities to recreate RS and TOR firing properties in simulations. In the model, the recorded whole-cell I_SA_ potassium currents and firing properties were best represented if RS cells were equipped with a single type of I_SA_ potassium conductance (I_SA_RS). Whereas, I_SA_ currents of TOR cells were reproduced by a mixture of two I_SA_ potassium conductances, including I_SA_RS and a left-shifted, slowly inactivating current (I_SA_TOR) in a 3:1 ratio. This mixed I_SA_ (I_SA_TOR+RS) is not only consistent with the variability of the patch current kinetics but also reproduces the properties of whole-cell currents (Figure 4B). After adding I_SA_RS or I_SA_TOR+RS to the core CCK+IN properties (Bezaire et al., 2016) in current-clamp simulations, the models reproduced the common firing properties of CCK+INs (including AP width, peak and frequency and AHP shape), but in the presence of I_SA_TOR+RS all five cells showed MP-dependent firing with similar temporal dynamics and voltage dependence as recorded experimentally. When the same cells were equipped with I_SA_RS only, they turned to RS firing type (Figure 4 – figure supplement 1E-F). Thus, exchange of I_SA_RS to I_SA_TOR+RS alone is sufficient to generate TOR properties, even if the reconstructed cell originally belonged to the RS type and vice versa.

Surprisingly, these results also suggest that TOR firing required much less (57.5%) I_SA_ potassium conductance in total, than RS firing in the same reconstructed cells. This could be an additional consequence of the left-shifted activation of I_SA_ in TOR cells (further supports for this argument will follow below).

### TOR cells are selectively silenced by ISATOR in a narrow range of oscillatory states

Next, we simulated the activity of RS and TOR firing cells during *in vivo*-like oscillating network conditions using excitatory and inhibitory inputs arriving onto the somato-dendritic axis of the five models of CCK+INs. The occurrence of excitatory events was clustered and tuned to frequencies ranging from 1 to 100 Hz and their kinetics were deducted from recordings from CCK+INs (see Methods). All five cells received the same input patterns and the strengths of excitation was varied (by changing the number of EPSCs, Figure 4C), which resulted in a wide range of spiking frequencies in CCK+INs representing the frequency ranges that have been recorded *in vivo* (Klausberger et al., 2005, Lasztoczi et al., 2011).

Next, we compared the average spiking of I_SA_TOR+RS potassium conductance-equipped CCK+INs (n = 5 cells) with the spiking of the same cells during the same conditions except that they were equipped only with I_SA_RS. As Figure 4 panel D shows, in most conditions, the presence of I_SA_TOR+RS instead of I_SA_RS did not markedly reduce the firing rates. However, CCK+INs were efficiently silenced by I_SA_TOR+RS in 8-15 Hz input frequency regimes. On average, 39.1 ± 0.6% fewer APs were evoked (see red areas in the middle of the Figure 4D graph and example traces). In contrast, during lower and higher input regimes the presence of I_SA_TOR+RS reduced firing only slightly (spiking was decreased by 8 ± 0.3% and 7.5 ± 0.2%, between 1-6 Hz and 25-100 Hz).

Thus, the results of these simulations suggest that I_SA_TOR+RS conductance alone enables CCK+INs to be selectively silenced during 8-15 Hz input regime, which adds a novel level of complexity to the diverse functions of GABAergic cells and can contribute to their observed heterogeneous firing during different network states (Klausberger et al., 2005, Klausberger and Somogyi, 2008, Lasztoczi et al., 2011). The input frequency dependence of the inhibition of firing can be explained by the specific temporal properties of I_SA_TOR+RS. Specifically, in addition to the voltage dependence of activation and inactivation (Figure 3), the time constant of inactivation (Figure 4 – figure supplement 1D) and the recovery from inactivation determines the different availability of these currents during various oscillatory states. Thus, minor modifications in the properties of I_SA_ enabled distinct functions in individual cells that otherwise belong to the same neuronal class.

### Kv4 channels are responsible for both types of ISA currents in CCK+INs

Next, we investigated the identity of potassium channel subunits responsible for the differences of I_SA_ in RS and TOR cells and for the MP-dependent firing in TOR cells. Neither TEA (0.5 and 10 mM, blocking Kv3 channels) nor low concentration of 4-AP (100 µM, blocking Kv1 and Kv3 channels) eliminated the initial firing gap in TOR cells (Figure 5 – figure supplement 1). Only a high concentration of 4-AP (5 mM) was able to diminish the TOR phenomenon in CCK+INs. These results, together with the low voltage activation properties, suggest a key role for Kv4 channels (Lien et al., 2002). Kv4.3 has been shown in hippocampal CCK+INs (Bourdeau et al., 2007, Kollo et al., 2006). To specifically test the contribution of Kv4 channels to the TOR phenomenon, we applied Heteropodatoxin-1 (HpTX, 1 µM), which selectively, but only partially, blocks Kv4.2 and Kv4.3 subunit-containing channels (DeSimone et al., 2011, Sanguinetti et al., 1997). In the presence of HpTX, significantly more APs were evoked in TOR cells during the first 125 ms of the stimulus compared to control conditions in the same cells before HpTX application (Figure 5A, from -80 mV preceding MP, 9.7 ± 2.9% vs 5.1 ± 2.2%, respectively, *p=0.0037, t(5)=-5.12, paired t-test*). Furthermore, the delay of the first APs was reversibly shortened from 239 ± 61 ms to 116 ± 38 ms (Figure 5B, *p = 0.01, t(7)=3.493, paired t-test, n = 8 TOR cells*). However, HpTX did not change the number and temporal distribution of APs in TOR cells from -60 mV (data not shown), nor the firing of RS cells neither at -80 nor -60 mV. HpTX did not influence the half-width of the APs (TOR cells: control: 0.48 ± 0.02 ms, HpTX: 0.52 ± 0.03 ms, *p = 0.15385, t(7) = -1.59901*, n = 8; RS cells: control: 0.55 ± 0.04 ms HpTX: 0.59 ± 0.06 ms, *p = 0.11016, t(7) = -1.82865*, n = 8). Thus, HpTX-sensitive, Kv4-mediated currents are crucial for the TOR phenomenon.

**Figure 5.**
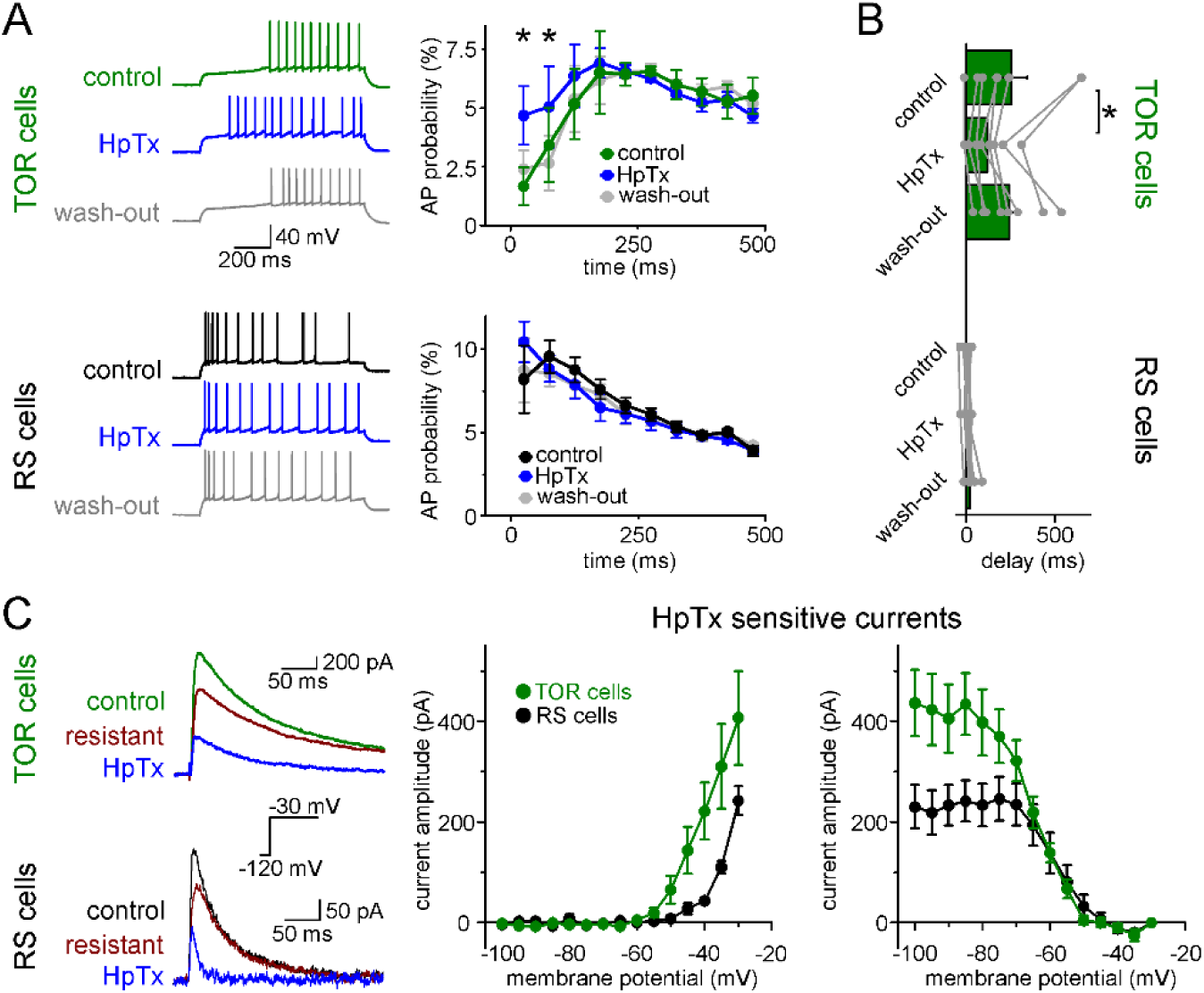
Blockade of Kv4 channels selectively inhibits TOR phenomenon. **A**. Representative CCK+INs firing patterns recorded before, during and after HpTx application (in response to identical current steps) and the average AP probabilities in TOR (n = 6) and RS cells (n = 9) from hyperpolarized MP ranges (−82 to -77 mV). **B**. Average delay of the first AP and reversible effect of HpTX. Connected symbols represent individual measurements (*paired t-test: p = 0.01008*). **C**. HpTx-sensitive I_SA_ in representative TOR and RS cells (left) and the average voltage dependence (n = 7 for both TOR and RS cells).

**Figure 5-figure supplement 1.**
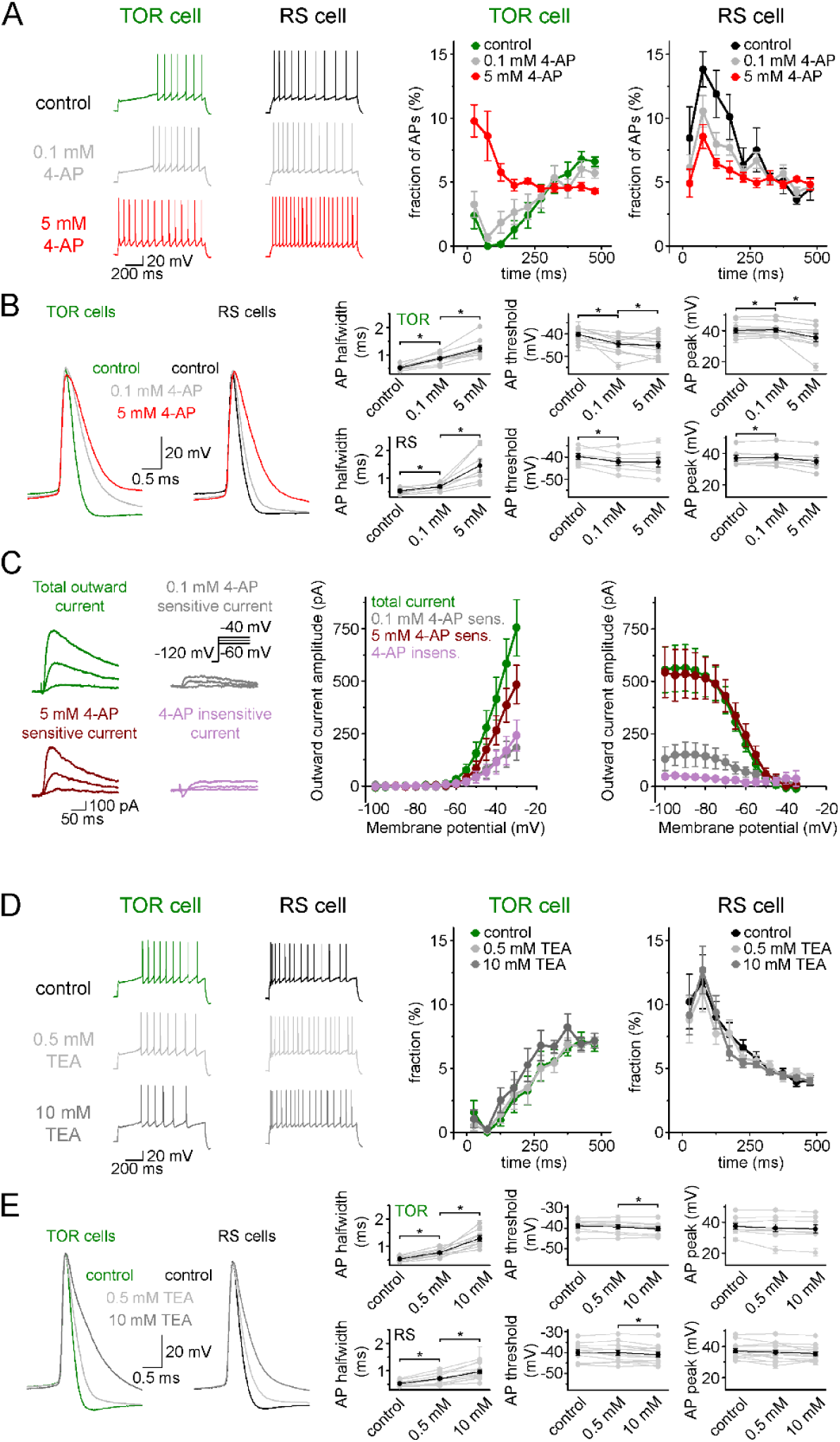
TOR phenomenon is not affected when Kv1 and Kv3 channels are inhibited. **A**. Inhibition of spiking in TOR cells was eliminated by only a high concentration of 4-aminopyridine (4-AP, 5 mM, red traces), but not by low concentration (0.1 mM, grey traces). **B**. As expected, both low and high concentrations of 4-AP affect AP width and threshold due to blockade of various potassium channels (grey symbols represent individual experiments, black symbols show average results). **C**. Representative traces on the left show control and 4-AP-sensitive and insensitive currents from a TOR cell and the graphs show the average voltage dependence of these components (n = 5 TOR cells). Majority (64.2% at -30 mV) of outward currents in TOR cells are sensitive to high, but not to low concentration 4-AP. **D**. Firing patterns of TOR and RS cells are not affected by low (0.5 mM, light grey) and high concentrations (10 mM, dark grey) of tetraethylammonium (TEA, n = 9 TOR and n = 13 RS cells). **E**. The effect of TEA on AP shape of TOR and RS cells (*: p < 0.05, *paired sample t-test*).

Next, we analyzed the contribution of HpTX-sensitive currents to I_SA_ in TOR and RS cells. Both types had substantial amounts of HpTX-sensitive currents. However, their properties were different in TOR and RS cells similarly as seen for their total I_SA_. In TOR cells, the HpTX-sensitive current activated at more negative voltage than in RS cells (Figure 5C). The time constant of inactivation of the HpTX-sensitive currents was slower in TOR cells than in RS cells (49.1 ± 12.6 ms, and 11.8 ± 1.6 ms, respectively, at -25 mV). The voltage dependence of inactivation of the HpTX-sensitive current was also left-shifted in TOR relative to RS cells (V_1/2_ TOR: -64.0 ± 0.6 mV, RS: -56.8 ± 1.3 mV). Considering the similar kinetics and voltage dependence of total and HpTX-sensitive I_SA_ currents and the partial Kv4-blocking ability of HpTX (DeSimone et al., 2011, Sanguinetti et al., 1997), these results suggest that the majority of inactivating I_SA_ potassium currents in both types of CCK+INs are mediated by Kv4 channels.

Next, we tested the presence of Kv4 channels in RS and TOR cells using the above SC-RNAseq data (Figure 6A-B). Kv4.2 was absent in most of the tested 17 CCK+INs confirming previous findings (Bourdeau et al., 2007, Rhodes et al., 2004). The mRNA of Kv4.3 subunits was detected in most CCK+INs (16 out of 17 cells), including both RS and TOR types, in agreement with the findings that both RS and TOR cells have HpTX-sensitive currents. Interestingly, in line with the predictions of the simulation, the average RNA copy number of Kv4.3 was slightly higher in the tested RS cells compared to TOR cells. Note that the SC-RNAseq data revealed differences in additional Kv channels between RS and TOR cells such as Kv3.2, Kv1.3 and Kv1.6 subunits (Figure 6A). Because these subunits do not generate low-voltage-activated, inactivating currents, we did not perform experiments to investigate the currents generated by these subunits (Lien et al., 2002, Figure 5 – figure supplement 1). We next performed immunohistochemistry to localize Kv4.3 subunits in biocytin-filled TOR and RS cells. In general, we observed an intense neuropil labelling for Kv4.3 in DG, CA3 strata radiatum and oriens, but not in the CA1 area. A subset of INs were also labelled throughout the hippocampus (Bourdeau et al., 2007, Rhodes et al., 2004). Kv4.3 proteins appeared to be enriched in the somatic and dendritic plasma membranes, but the subcellular distribution was often uneven and clustered (Figure 6D and Figure 6 – figure supplement 1A) in line with the known distribution pattern of the Kv4.3 subunit (Kollo et al., 2006). In agreement with the pharmacological and SC-RNAseq, data we detected Kv4.3 proteins (Figure 6B, C) in both TOR (15 out of 20 tested cells) and RS CCK+INs (19 out of 23 cells). However, the immunosignal for Kv4.3 was usually stronger in RS than in TOR cells even within the same sections. This tendency was also seen in the CA3 area obtained from perfusion-fixed brain (Figure 6 – figure supplement 1A), where Kv4.3 signal was detectable at strong or moderate level in most CCK/Satb1-cells (putative RS cells, 99%, 338 out of 342 tested cells), whereas the labelling was weak or hardly detectable in most CCK/Satb1+ cells (putative TOR cells, 97%, 78 out of 80 tested cells). TOR and RS cells not only possess Kv4.3-mediated I_SA_ with different activation and inactivation kinetics, but TOR cells seem to achieve larger charge transfer via lower density of Kv4.3 channels. What could mediate such differences?

**Figure 6.**
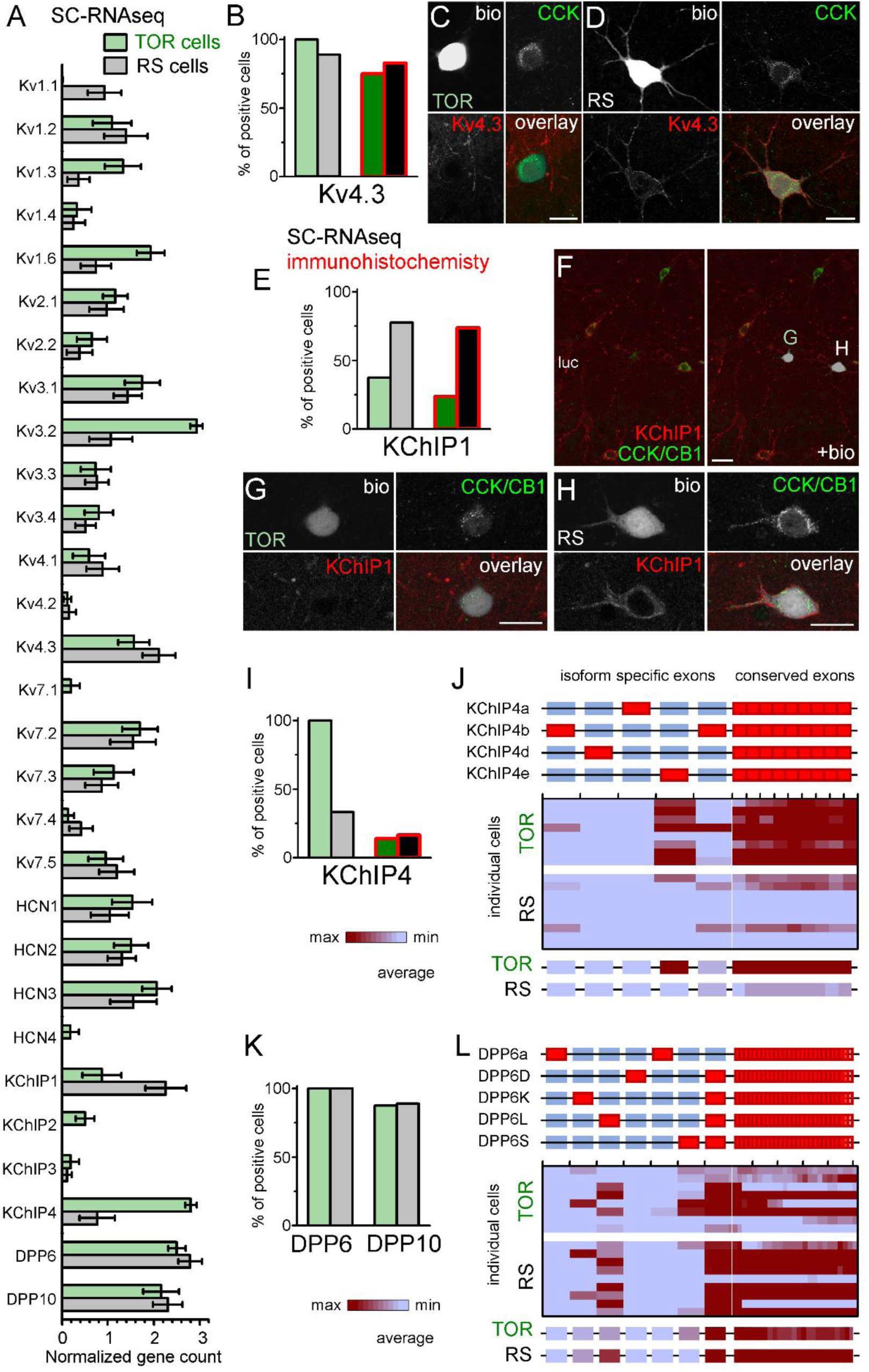
Similar Kv4.3-expression and different auxiliary subunit, KChIP and DPP-content in CCK+INs. A. Normalized gene count of primary and auxiliary subunits of voltage-gated potassium channels from single cell RNAseq data of TOR (n = 8) and RS (n = 9) cells. **B**. Percentage of recorded cells with detectable levels of Kv4.3 mRNA (left bars, n = 8 and 9 cells) and protein (right bars, n = 20 and 23 tested cells). **C-D**. Immunofluorescent co-localization of CCK and Kv4.3 in a TOR **(C)**, and a RS cell **(D)** in CA3 stratum lucidum. **E**. Percentage of recorded cells with detectable levels of KChIP1 mRNA (n = 8 and 9 cells) and proteins (n = 20 and 27 cells). **F-H**. Immunofluorescent co-localization of CCK with CB1 (green) and KChIP1 in a TOR **(G)**, and a RS cell **(H)** from the same slice shown in low magnification image **(F). I**. Percentage of recorded cells with detectable levels of KChIP4 mRNA (left bars, n = 8 and 9 cells) and protein (right bars, n = 29 and 30 tested cells). **J**. Major KChIP4 splicing isoforms consist of different exons in the N-terminal region (represented as red boxes). Each row represents a single cell in the color-mapped data and columns correspond to individual exons aligned to the schematic illustration of isoforms above. Red and blue colors code high and low mRNA levels, respectively. The average exon counts from the two types of CCK+INs (n = 8 and 9) are shown at the bottom using the same color code scheme. **K**. Percentage of recorded cells with detectable levels of DDP6 and DPP10 mRNA. **L**. Assembly of major DPP6 isoforms and exon levels in individual CCK+INs are shown as above (J).

**Figure 6 – figure supplement 1.**
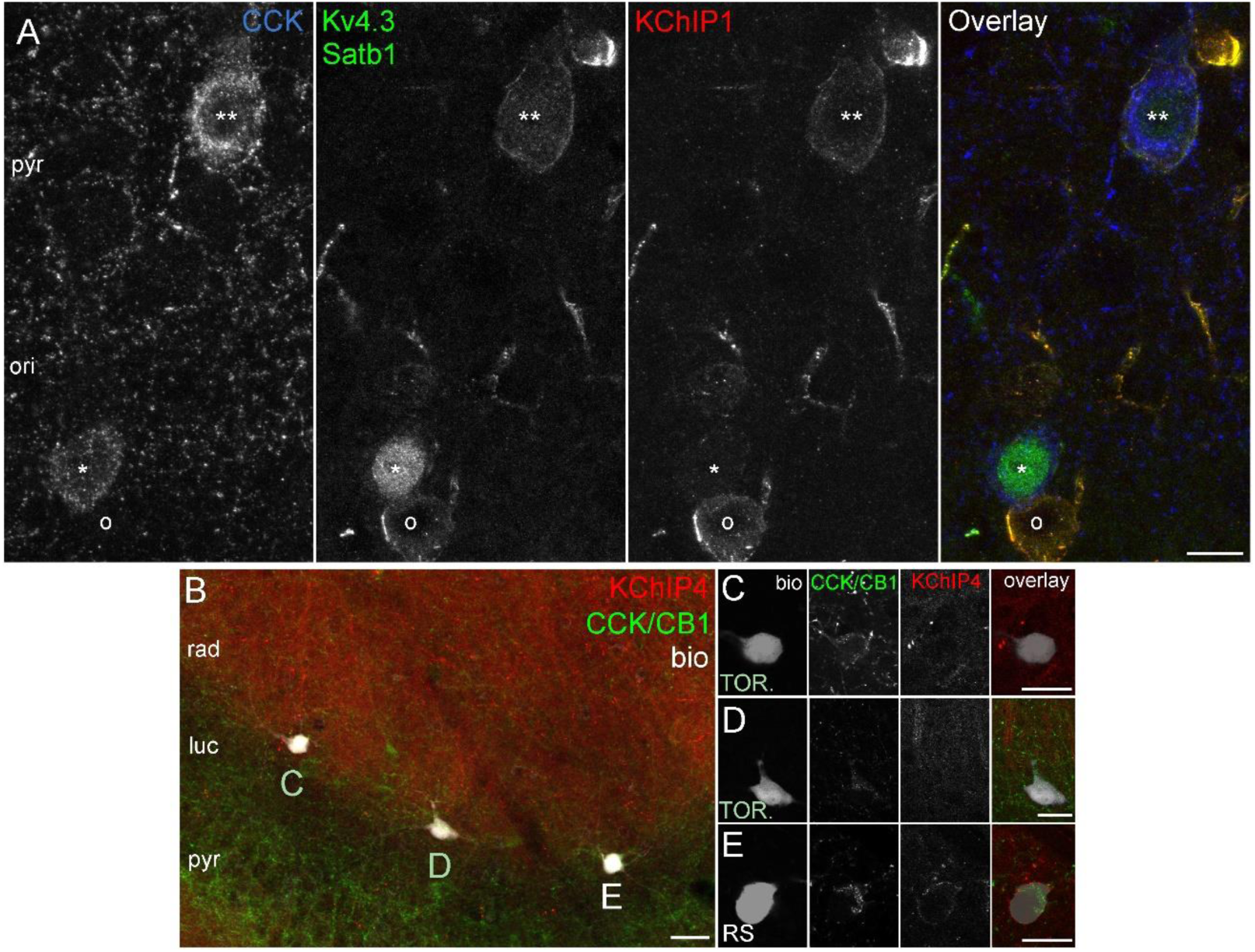
Kv4.3 and KChIP immunohistochemistry in perfusion or immersion fixed CA3 slices. **A**. CCK+ cells in CA3 obtained from perfusion fixed brain. One CCK+ cell (**) is outlined by intense Kv4.3 and KChIP1 immunolabelling. In the other CCK+ cell (*) that is strongly labelled for Satb1 in the nucleus, the Kv4.3 and KChIP1 signals are only weakly detectable, whereas they both intensely decorate a neighbouring CCK-negative neuron (o). **B**. Low magnification image showing simultaneous immunolabelling of KChIP4 (red) and CCK/CB1 (green) in an acute CA3 slice, in which a RS cell and two TOR cells were filled with biocytin (left) at the border of strata radiatum/lucidum in CA3a. Intense KChIP4 immunolabelling is present in the neuropil of stratum radiatum mainly associated with dendrites of pyramidal cells (Rhodes et al., 2004). **C-E**. High magnification images showing an enlarged view of the biocytin filled cells shown in (**B**). KChIP4 is not detectable in the two TOR cells (**C, D**), but its weak presence was detected in the neghbouring the RS cell (**E**), but. Scale bars: 10 µm (**A**) 20 µm (**B-E**).

### Auxiliary subunits of Kv channels in TOR and RS cells

Core Kv4 channel proteins form ternary complexes with dipeptidyl aminopeptidase-like proteins (DPLPs, including DPP6 or DPP10) and K^+^ channel interacting proteins (KChIP1-4 from Kcnip1-4 genes), which fundamentally change the current properties. The modulatory effects are highly variable and depend on the type and splice isoforms of the available auxiliary subunit. For example, most KChIP isoforms enhance surface expression and accelerate inactivation and recovery from inactivation (Jerng and Pfaffinger, 2014, Pongs and Schwarz, 2010). However, the so-called transmembrane KChIPs (tmKChIPs) have opposite effects as they can retain Kv4 from the plasmamembrane and decelerate inactivation (Holmqvist et al., 2002, Jerng and Pfaffinger, 2008, Jerng and Pfaffinger, 2014, Pruunsild and Timmusk, 2012).

KChIP1, as one of the classical KChIPs, accelerates inactivation and increases surface expression of Kv4.3 channels (Beck et al., 2002, Bourdeau et al., 2011, Jerng and Pfaffinger, 2014, Pongs and Schwarz, 2010). We detected KChIP1 mRNA in most RS cells (7 of 9 tested), but it was absent in 5 out of 8 TOR. The average mRNA level was significantly higher in RS than in TOR cells (2.26 ± 0.44, TOR: 0.87 ± 0.42, Mann-Whitney, p = 0.013, z = -2.475). Detection of the KChIP1 protein by immunohistochemistry revealed an even clearer distinction between RS and TOR cells (Figure 6E). The KChIP1 protein was detected in the majority of tested RS cells (20 out of 27), whereas only few TOR cells showed positive immunoreaction (5 out of 21) and the signal in these cells was weaker than in RS cells (Figure 6F-H). KChIP1 signal was present not only in the plasma membrane but also in the cytosol, in agreement with their trafficking role (Pongs and Schwarz, 2010). This data from recorded cells were confirmed by the analysis of CCK+INs in perfusion fixed brains (Figure 6 - figure supplement 1A). We detected strong KChIP1 signal in the majority of CCK+/Satb1-cells (corresponding to RS cells, 122 out of 132 tested cells, 92.4%). In contrast, only 4.9% of CCK+/Satb1+ cells (corresponding to TOR cells, 2 out of 41 tested cells) showed strong KChIP1 immunosignal. We did not detect significant amounts of KChIP2 and KChIP3 mRNAs in CCK+INs.

The universal mRNA sequence of KChIP4s was detected in all TOR cells (8 out of 8), whereas it occurred in 3 out of 9 RS cells. The available antibody detected KChIP4 protein only in a very few CCK+INs regardless of their firing type (4 out of 29 TOR, 5 out of 30 RS cells, Figure 6I and Figure 6 – figure supplement 1B-E), thus, our observation with immunohistochemistry apparently contradicts the mRNA data. In general, the relatively weak KChIP4 immunosignal in CCK+INs was surrounded by strong neuropil labelling in the stratum radiatum (Figure 6 - figure supplement 1B). While KChIP1 is expressed only in INs, KChIP4 is known to be associated with Kv4.2 channels in pyramidal cells as well (Rhodes et al., 2004). This is in accordance with our observation of KChIP4 immunosignal at high magnification (Figure 6 - figure supplement 1D), where it appears associated with tube-like structures, likely representing the plasma membranes of putative pyramidal cell dendrites. However, our KChIP4-specific antibody was raised against a long amino acid segment, which includes the highly variable N-terminal region, which endow the various KChIP4-isoforms with different effects on Kv4 channel function (Holmqvist et al., 2002, Jerng and Pfaffinger, 2008, Jerng and Pfaffinger, 2014, Pruunsild and Timmusk, 2012). Therefore, we further analyzed the mRNA data at the level of individual KChIP4 exons (Figure 6J). We detected the KChIP4e isoform-specific exon in 7 TOR cells (out of 8 tested). KChIP4e belongs to the so-called transmembrane KChIPs (tmKChIPs) (Jerng and Pfaffinger, 2008), which, in contrast to most KChIP types and isoforms, do not promote the plasma membrane expression of Kv4 and have opposite effects on the inactivation kinetics (Jerng and Pfaffinger, 2008, Pruunsild and Timmusk, 2012). KChIP4b, which acts as classical KChIPs, was detected in only one TOR cell (from 8 tested). KChIP4e specific exons were detected only in one RS cell. The two other RS cells that also had a small amount of KChIP4, expressed the KChIP4b isoform. Altogether, these data indicate that RS cells express KChIP1, whereas TOR cells primarily express KChIP4e. We suggest that the differential influences of these auxiliary subunits underlie some of the distinct properties of I_SA_ in RS and TOR cells because KChIP1 is known to accelerate inactivation and increase surface expression, whereas tmKChIPs, such as KChIP4e, do not facilitate Kv4 plasma membrane trafficking and confer slow channel inactivation.

KChIPs alone do not account for all differences between I_SA_ currents in TOR and RS cells, including differences in voltage-dependence. DPP10 and DPP6 proteins effectively shift the voltage dependence of Kv channel activation and inactivation (Jerng et al., 2007, Jerng and Pfaffinger, 2012, Jerng and Pfaffinger, 2014, Nadal et al., 2006, Nadal et al., 2003, Pongs and Schwarz, 2010). Therefore, next we analyzed the expression of these molecules at the mRNA level (Figure 6K-L). DPP10 (conserved segments) was present in both TOR and RS cells (7 out of 8, and 8 out of 9 cells, respectively). The DPP10c isoform was detected in significant amounts in all cells regardless of their firing type. DPP6 was also detected in most CCK+INs from both types. The primary isoform in RS cells was the DPP6L. In contrast, DPP6S isoform occurred in most TOR cells (7 out of 8 cells) and DPP6L-specific exons were also detected in several cells (4 out of 8 TOR cells). Thus, in addition to KChIPs, expression of DPLP-isoforms correlates with the firing types of CCK+INs and their combined modulatory effects may underlie the different properties of Kv4.3-mediated I_SA_ currents, which is responsible for their different excitability.

## Discussion

CCK+INs can dynamically select cell assemblies and control their activity during theta oscillations, which dominate hippocampal operation during spatial navigation and memory tasks (Klausberger and Somogyi, 2008). Therefore, it is particularly important that the excitability of TOR and RS cells are drastically different when they are excited within 8-15 Hz input range. This suggests that under basal conditions it is unlikely that TOR cells can substantially contribute to theta oscillations unless they are primed by preceding depolarizations. By contrast, TOR and RS cells may play similar roles during lower and faster network oscillations. Thus, because distinct network state-dependent contribution is considered as a cell type classification criteria (Klausberger and Somogyi, 2008), TOR and RS cells can be viewed as separate partitions within the CCK+IN class. Furthermore, the TOR phenomenon can also contribute to the variable recruitment of individual CCK+INs during subsequent theta cycles (Freund and Katona, 2007, Klausberger et al., 2005).

Despite the functional distinction between TOR and RS cells, their morphology, basic electrophysiological characteristics and most genes are surprisingly similar. One notable difference is the presence of transcription regulator Satb1 in most TOR cells and absence in most RS cells. However, we did not detect systematic differences in the complete transcriptome of TOR and RS cells. Even those genes and proteins were present in both types similarly that have been shown previously to delineate certain subpopulations within CCK+INs (such as VGluT3 or Calb1). Thus, TOR and RS cells cannot be distinguished based on their general gene expression profiles, which is recently used for identification of individual cell types (Fuzik et al., 2016, Harris et al., 2018). Furthermore, the morphology of RS and TOR cells covers the same diversity including soma and dendrite targeting axons and we did not recognize differences in their AP shape and synaptic inputs. Thus, it is difficult to determine whether TOR and RS CCK+INs comply with conventional definitions of cell types in spite of their potential different contributions to network operations.

### Same channel protein but distinct auxiliary subunits are responsible for the different I_SA_ currents and for the different functionality of TOR and RS cells

Our simulations showed that Kv4.3 current alone is sufficient for the oscillation frequency-dependent functionality among CCK+INs because changing only I_SA_RS to I_SA_TOR+RS in the same realistically modelled CCK+INs specifically silenced them during 8-15 Hz input regimes and reproduced the recorded differences in their firing. The recordings confirmed the crucial role of Kv4.3 channels in TOR firing. However, RS cells also express Kv4.3 channels, and in fact, their density is higher in RS cells compared to TOR cells. Thus, paradoxically, in spite of Kv4.3 mediated currents are responsible for the unique and distinctive firing properties of TOR cells, Kv4.3 currents are present in both cell firing types. We found the potential explanation that solves this paradox in differences of the auxiliary subunits that can modify the function of Kv4.3 channels.

KChIP1 is strongly expressed by RS cells, whereas most TOR cells lacked this cytosolic auxiliary subunit. The known effects of the KChIP1 (Beck et al., 2002, Bourdeau et al., 2011, Jerng and Pfaffinger, 2014, Pongs and Schwarz, 2010) correlate well with the properties of the I_SA_ in RS cells. As a classical KChIP, KChIP1 increases surface expression of Kv4 channel complex. Indeed, we measured larger density of I_SA_ current in outside out patch from RS cells than in TOR cells. We also observed that Kv4.3 immunosignal was usually stronger in RS cells than in TOR cells. Furthermore, we were able to reproduce TOR and RS firing phenotypes in the realistically simulated conditions only if the total amount of I_SA_ conductance was larger in RS cells than in TOR cells. The apparent paradox between larger current density and the smaller measured current amplitude near the AP threshold in RS cells derives from the differences in the voltage dependence of activation of I_SA_ in RS and TOR cells. I_SA_ channels are submaximally activated at physiological subthreshold voltage. However, because of the left-shifted activation curve, a much larger fraction of I_SA_ channels are activated in TOR cells at lower voltage ranges. Therefore, larger currents can be generated even by a smaller number of channels. Another substantial influence of KChIP1 on Kv4.3 is the acceleration of steady-state inactivation kinetics and the recovery time from inactivation (Beck et al., 2002, Bourdeau et al., 2011, Jerng and Pfaffinger, 2014, Pongs and Schwarz, 2010). These are also correlated well with the differences of I_SA_ in RS and TOR cells, as both parameters were much faster in the former type of cells.

KChIP4 was the dominant KChIP in TOR cells and it was not detected in most RS cells. Our data also showed that TOR cells expressed a special splicing isoform, KChIP4e, which belongs to the so-called tmKChIPs (Holmqvist et al., 2002, Jerng and Pfaffinger, 2008, Jerng and Pfaffinger, 2014, Pongs and Schwarz, 2010). In contrast to classical KChIPs, all tmKChIPs including KChIP4e do not facilitate surface expression of Kv4 channels (Holmqvist et al., 2002, Jerng and Pfaffinger, 2008, Pruunsild and Timmusk, 2012). As explained above, several lines of evidence suggest lower Kv4.3 channel density in the KChIP4e-expressing TOR compared to KChIP1-expressing RS cells. The presence of tmKChIPs results in slow inactivation kinetics, often slower than that of the solitary Kv4 channels (Holmqvist et al., 2002, Jerng et al., 2007, Jerng and Pfaffinger, 2008, Tang et al., 2014). As a consequence of enhanced closed-state inactivation of Kv4.3 channels, KChIP4a causes a leftward shift in the voltage dependence of inactivation (Tang et al., 2013, Tang et al., 2014). Albeit it is a likely possibility due to the similarity of their N-terminal domains, it remains to be answered whether the effects of KChIP4e on Kv4.3 kinetics are similar to the other studied tmKChIPs. The slower inactivation and recovery of I_SA_ in KChIP4e-expressing TOR cells were critical for their unique functionality during 8-15 Hz network states because they define the availability of the I_SA_ conductance. KChIP4 was present only in three RS cells and two of them expressed the KChIP4b isoform, which is a classical KChIP that, similarly to KChIP1s results in much faster inactivation and recovery (Jerng and Pfaffinger, 2008). We should also note that the apparent discrepancy in KChIP4 detection between the mRNA and protein levels may also be explained by the presence of different isoforms, as our antibody may have targeted the highly variable N-terminal region.

Both DPLPs were present in TOR and RS cells. DPP10c isoform, which is known to affect voltage dependence of Kv4 channels, but does not accelerate inactivation (Jerng et al., 2007), was ubiquitous in both types of CCK+INs. Albeit all tested CCK+INs had a significant amount of DPP6 mRNA, TOR and RS cells expressed different isoforms. RS cells expressed only DPP6L and in TOR cells the primary isoform was DPP6S (but DPP6L was also present in several cells). DPP6S can contribute to the left-shifted voltage dependence of activation of I_SA_ in TOR cells. The left-shifted voltage dependence is important for the sufficient prevention of spiking. The dual set of DPP6 proteins fits well with our observations that many properties of TOR cells can be described only if two populations of Kv4.3-mediated currents are present. Altogether, these observations suggest that apparently small modifications in the available components of ion channel complexes underlie the different functions of TOR and RS cells.

The net effects of KChIPs and DPLPS are not simply the sum of the effects of individual subunits. Some subunits either dominate others or the interaction can result in unique combinations of effects (Jerng et al., 2005, Nadal et al., 2006, Zhou et al., 2015). For example, when DPP6K is present, KChIP4e causes a leftward shift of the voltage dependence of inactivation and deceleration of the recovery from inactivation compared to other KChIPs (Jerng and Pfaffinger, 2012). In this regard, it is an important observation for our results that DPP6S, unlike some other DPLPs, cannot overcome the tmKChIP4-mediated deceleration of inactivation (Jerng et al., 2007, Jerng and Pfaffinger, 2008, Seikel and Trimmer, 2009). Thus, the expression of DPP6S in TOR cells may preserve the unique modulatory effects of KChIP4e. Because of the composition of these ternary channel complexes by multiple proteins and splicing variants, whose interactions are not yet characterized, the exact contribution of individual components of DPP6S/L/DPP10c-KChIP4e-Kv4.3- and DPP6L/DPP10c-KChIP1-Kv4.3-complexes cannot be predicted yet. Nevertheless, the higher channel density, faster inactivation kinetics and faster recovery from inactivation of I_SA_ in RS compared to TOR cells are consistent with the differential expression of KChIP1 and KChIP4e, and the cell type-specific expression and contributions of DPP10c, DPP6L and DPP6S are expected to explain unique voltage-dependency in the two types of CCK+INs without the involvement of additional differences between RS and TOR cells. Thus, the different firing properties and responsiveness during 8-15 Hz network states of RS and TOR cells are established by surprisingly minor modifications in a few auxiliary subunits.

## Methods

Animal protocols and husbandry practices were approved by the Institute of Experimental Medicine Protection of Research Subjects Committee (MÁB-7/2016 for slice recording and anatomy experiments and MÁB-2/2017 for immunolabelling experiments in perfusion fixed brains). by the Veterinary Office of Zurich Kanton (ZH241/15, single cell RNAseq experiments).

### Slice preparation, solutions and chemicals

Hippocampal slices were prepared from 21-33 days old Wistar rats (deeply anaesthetized with isoflurane) in ice-cold artificial cerebrospinal fluid (85 mM NaCl, 75 mM sucrose, 2.5 mM KCl, 25 mM glucose, 1.25 mM NaH_2_PO_4_, 4 mM MgCl_2_, 0.5 mM CaCl_2_, and 24 mM NaHCO_3_, Leica VT1200 vibratome). The slices were incubated at 32 °C for 60 minutes after sectioning and were then stored at room temperature until they were used. The normal recording solution was composed of 126 mM NaCl, 2.5 mM KCl, 26 mM NaHCO_3_, 2 mM CaCl_2_, 2 mM MgCl_2_, 1.25 mM NaH_2_PO_4_, and 10 mM glucose. For standard recordings, pipettes were filled with an intracellular solution containing 90mM K-gluconate, 43.5 mM KCl, 1.8 mM NaCl, 1.7 mM MgCl_2_, 0.05 mM EGTA, 10 mM HEPES, 2 mM Mg-ATP, 0.4 mM Na_2_-GTP, 10 mM phosphocreatine, and 8 mM biocytin (pH 7.2; 270–290 mOsm). Chemicals for the intra- and extracellular solutions were purchased from Sigma-Aldrich, ion channel blockers were from Tocris or Alomone and fluorophores were from Invitrogen.

### Somatic recordings

For recordings, cells in slices were visualized with an upright microscope (Eclipse FN-1; Nikon) with infrared (900 nm) Nomarksi DIC optics (Nikon 40x NIR Apo N2 NA0.8W objective). Electrophysiological recordings were performed at 34.5-36°C. During current-clamp recordings, firing patterns were elicited by square shaped current pulses with increasing amplitudes (starting from -100 pA up to 700 pA, Δ20 pA, duration: 1 sec) or with standard steps (eliciting average firing between 10 -20 Hz), which was preceded by 3-second-long different amplitude holding current steps () to reach preceding MP range between -90 and - 50 mV. The action potential (AP) distributions were calculated from all recorded traces, which contained action potentials and binned by 50 ms from each recorded cell. In these recordings, the pipette capacitance was neutralized (2.5-5 pF remaining capacitance) and bridge balance compensation was set to eliminate apparent voltage offsets upon current steps. Voltage values were not corrected for the liquid junction potential (theoretically: -15.4mV). Traces were low pass filtered at 6–20 kHz and digitized at 40–100 kHz using a Digidata 1440A interface (Molecular Devices).

For voltage-clamp recordings, cells were patched in normal extracellular solution to first record their firing patterns, and then 2.5 µM TTX and 10 µM ZD7288 was added to block sodium and Ih currents respectively. Voltage protocols for potassium current measurements consisted of a 300 ms long conditioning pulse at -120 mV, followed by a 300 ms long voltage steps between -120 and -20 mV (for current activation and decay time constant measurements), and a last voltage step to -30 mV for 100 ms (to measure voltage dependence of potassium current inactivation). Series resistances were between 6-20 MΩ (75-80% compensated with 53 µs lag) and were constantly monitored. Data were discarded if the series resistance changed more than 25%.

The recovery from inactivation test protocol consisted four voltage steps: (1) -120 mV for 345 ms, allowing full recovery, (2) -30 mV for 500 ms, allowing complete inactivation, (3) -65, -75, -85 or -120 mV steps with variable duration between 1-233 ms, (4) -30 mV test pulse, where area of current was analyzed according to the voltage and duration of the preceding recovery step. Control traces (whose duration was identical to test pulses and steps 1 and 3 were set to -50 mV) were subtracted from each traces allowing the isolation of _ISA_ currents. Furthermore, these measurements were performed in the presence of TTX, 1 mM 4-AP and 10 mM TEA to reduce contamination from non-Kv4 channels.

### Outside-out patch recording

To obtain outside-out patch recordings, first we made somatic whole-cell recordings from the selected cells using IR-DIC optics. This was necessary for classifying their firing as TOR or RS cells, for loading biocytin for subsequent CCK immunolabelling, and for loading 20 µM Alexa-594 fluorescent dye, which allowed the visualization of their dendrites. After at least a 5 -minute-long loading period the somatic pipette was retracted. Intact dendrites (30-80 µm from the surface) were visualized and patched using epifluorescent illumination (less than a minute illumination time). The thick-wall pipettes (resistance: 20-55 MΩ) were filled with 5 µM Alexa-594. After break-in, we confirmed that no neighboring structure was loaded with the fluorescent dye and outside-out configuration was achieved by the slow retraction of the pipette. By applying similar voltage steps as described for whole-cell voltage-clamp configurations, we recorded dendritic potassium currents (without pharmacological isolation). After outside-out patch recordings, the pipette was pushed into a Sylgard ball (Sylgard 184, Merck), and capacitive responses to voltage steps were recorded and subtracted from the capacitive responses recorded in outside-out patch configuration. The surface area of the patch membrane was calculated as described earlier (Gentet et al., 2000) with the specific membrane capacitance determined from 18 nucleated patch experiments (cm = 1.015 ± 0.014 µF/cm^2^). Distances of the recording sites from the soma were measured based on posthoc epifluorescent or confocal images. Somatic outside-out patches were obtained using the same protocol, with similar pipettes. Current traces were low pass filtered at 10 kHz and digitized at 100 kHz. Capacitive membrane responses were digitized at 250 kHz, without filtering. Leak and capacitive current components were subtracted during potassium current recordings using online P/–4 method. Inactivating potassium currents (at 0 mV) were isolated offline by subtracting currents obtained with a -50 mV prepulse from currents measured with a prepulse of -80 mV.

Anatomical and immunohistochemical characterization of CCK+ cells All recorded cells were filled with biocytin and processed for immunhistochemistry. After the recordings, slices were fixed in 0.1 M phosphate buffer containing 2% paraformaldehyde and 0.1% picric acid at 4 °C overnight. After fixation, slices were re-sectioned at 60 μm thickness (Neubrandt et al., 2018). Immunopositivity for CCK was tested with a primary antibody raised against cholecystokinin (1:1000, CCK, Sigma-Aldrich, Cat# C2581, RRID:AB_258806, rabbit polyclonal). Biocytin labeling was visualized with either Alexa 350-, Alexa 488-, Alexa 594- or Alexa 647-conjugated streptavidin. For additional neurochemical markers, further immunolabeling was used either against Vesicular Glutamate Transporter 3 (1:2000, VGluT3; Merck, Cat#AB-5421, RRID:AB_2187832; guinea pig polyclonal), Cannabinoid Receptor type 1 (1:1000, CB1R, Cayman Cat#10006590, RRID:AB_10098690, rabbit polyclonal), special AT-rich sequence-binding protein-1 (1:400, Satb1; Santa-cruz Cat#sc-5989, RRID:AB_2184337, goat polyclonal, 1:400), Reelin (1:400, Merck Cat#MAB5364, RRID:AB_2179313, mouse monoclonal) or calbindin (1:1000, Calb1; Swant Cat#300, RRID:AB_10000347, mouse monoclonal). Only those cells were included in the analysis, which were positive for CCK (or for CB1, n = 15 cells).

Known morphological subtypes of CCK+INs in the hippocampal CA3 region were determined based on the layer-preference of their axonal arborization. Schaffer collateral associated cells (SCA)(Cope et al., 2002) innervate dendrites in the stratum radiatum. Albeit the definition of basket cells (BCs) (Hendry and Jones, 1985) and mossy-fiber associated cells (MFA) (Vida and Frotscher, 2000) are clear, with termination zones in the strata pyramidale and lucidum, respectively, their practical identification is more complicated because both cell types have a substantial amount of axons in the adjacent strata, especially near to their soma. Therefore, only cells with extended axonal arbor (at least reaching 200 µm from soma) were identified either as BC or as MFA. In this distal region, BCs had clearly targeted stratum pyramidale, whereas distal axons of MFA ran in the stratum lucidum parallel with the cell layer, and they often invaded the hilar region of the dentate gyrus.

### Analysis of potassium channel subunits using immunohistochemistry

For perfusion fixed brain samples two (P25 and P45) Wistar rats were anesthetized and perfused through the aorta with 4 % paraformaldehyde and 15 v/v% picric acid in 0.1M Na-phosphate buffer (PB, pH = 7.3) for 15 min. Immunohistochemistry was performed on 70 μm thick free floating coronal sections from the hippocampus. We used a different fixation protocol for acute hippocampal slices containing biocytin-filled cells. After short recordings (5-15 minutes), slices were transferred to a fixative containing 2% paraformaldehyde and 15 v/v% picric acid in 0.1 M PB for 2 hours at room temperature. Slices were washed in PB, embedded in agarose and re-sectioned to 70-100 µm thick sections. Sections were blocked in 10% normal goat serum (NGS) in 0.5 M Tris buffered saline (TBS) for one hour and incubated in a mixture of primary antibodies for one overnight at RT. The following antibodies were used: rabbit polyclonal anti-CCK antibody (1:500, Sigma-Aldrich Cat# C2581, RRID:AB_258806) mixed either with a mouse monoclonal anti-KChiP1 (1:500, IgG1, UC Davis/NIH NeuroMab Facility Cat# 75-003, RRID:AB_10673162), a mouse monoclonal anti-KChiP4 (1:400, IgG2a, UC Davis/NIH NeuroMab Facility Cat# 75-406, RRID:AB_2493100), or a mouse monoclonal anti-Kv4.3 antibody (1:500, IgG1, UC Davis/NIH NeuroMab Facility Cat# 75-017, RRID:AB_2131966). In some sections, a rabbit polyclonal anti-CB1 receptor (1:2000, Cayman Chemical Cat# *10006590, RRID*:AB_10098690) was also used together with the CCK antibody to more reliably characterize filled cells. Potential TOR cells were identified by Satb1 immunolabelling in the perfusion fixed sections. Satb1 and Kv4.3 were visualized by the same secondary antibody but their labelling pattern could be reliably distinguished based on their different subcellular locations. The following secondary antibodies were used to visualize the immunoreaction: Alexa 488-conjugated goat anti-rabbit (1:500, Thermo Fisher Scientific), Cy5 conjugated goat anti-rabbit (1:500, Jackson ImmunoResearch), Alexa 488-conjugated goat anti-mouse IgG1 (1:500, Jackson ImmunoResearch), and Cy3 conjugated goat anti-mouse IgG1 or IgG2a (1:500, Jackson ImmunoResearch) IgG-subclass-specific secondary antibodies. Biocytin was visualized with Cy5 conjugated Streptavidin (1:1000, Jackson ImmunoResearch). High magnification fluorescent images were acquired with an Olympus FV1000 confocal microscope using a 60x objective (NA = 1. 35).

### Single-cell mRNA

Single-cell mRNA was performed using the Clontech’s SMARTer v4 Ultra Low Input RNA Kit. Cells were collected via pipette aspiration into sample collection buffer, were spun briefly, and were snap frozen on dry ice. Samples were stored at –80 °C until further processing, which was performed according to the manufacturer’s protocol. Library preparation was performed using Nextera XT DNA Sample Preparation Kit (Illumina) as described in the protocol. Then, cells were pooled and sequenced in an Illumina NextSeq500 instrument using 2 × 75 paired end reads on a NextSeq high-output kit (Illumina). After de-multiplexing the raw reads to single-cell datasets, we used Trimmomatic and flexbar to remove short reads, remove adapter sequences and trim poor reads. The remaining reads were aligned to the GRCm38 genome with STAR aligner. Aligned reads were converted to gene count using RSeQC. All data analyses were performed using python3. The analysis included the removal of poor quality cells (at least 3000 genes were detected in each cell), normalization of gene expression data using scran, and analysis of differential expression of genes across cell types. Sequences of splicing variants of KChIP4, DPP6, and DPP10 were validated in the NCBI Genebank and UCSC databases.

Genes related to apoptosis (Bcl2, Casp2, Casp8, Fas; not shown) were absent or present in low copy numbers in both RS and TOR cells indicating that the two firing phenotypes are not due to differential damage during slice preparation.

### Computational modelling

We performed computer simulations using the NEURON simulation environment (version 7.5 and 7.6, downloaded from http://neuron.yale.edu). To create a realistic model of CCK+INs, first we made detailed reconstructions of five biocytin labelled, electrophysiologically characterized cells (using Neurolucida and the Vaa3D software; Peng et al., 2010). The passive electrical parameters of the simulated cells were set as follows: first, axial resistance values were set to 120 Ωcm, then specific membrane capacitance and leak conductances (the substrate of membrane resistance) were fitted based on passive membrane responses to small amplitude (20 pA) current injections (ranges: cm: 1.05 – 0.82 µF/cm^2^, gl: 5*10^-5^ – 5.8*10^-5^ S/cm^2^). The set of active conductances were selected based on a previous publication (Bezaire et al., 2016). The properties (voltage dependence and kinetics) of these conductances were adjusted to reproduce firing characteristics representing our average measurements (AP parameters: AP peak, AP half-width, AHP amplitude, AP threshold, and firing parameters: accommodation of firing frequency, maximum AP frequency). Additionally, two variants of A-type potassium conductance models were created. First, the ratio of somatic and dendritic inactivating potassium conductances was set to 2.96:1 somatic to dendritic ratio (see Figure 4-figure supplement 1). Based on the observation, that patches pulled from RS cells produced inactivating potassium currents with similar MP-dependence to those recorded in whole-cell configuration, we modelled a single potassium conductance (I_SA_RS) constrained on somatic whole-cell recordings (HpTx-1 sensitive currents measured in RS cells, Figure 5C), with the appropriate uncompensated series resistance implemented in the models (2.76 ± 0.28 MΩ, n= 14). For TOR cell models we used a mixture I_SA_ consisting of I_SA_RS and I_SA_TOR, to account for the variability of patch currents from TOR cells. I_SA_TOR has a left shifted voltage dependence and slower inactivation than I_SA_RS. I_SA_TOR+RS reproduced potassium currents from our whole-cell measurements and the typical characteristics of TOR firing.

To investigate the behavior of these model cells in *in vivo* relevant conditions constructed physiologically plausible input conditions with a large number of temporally organized synaptic excitation and inhibition. For these, first we measured the amplitude and kinetics of glutamaterg and GABAerg currents in TOR and RS cells, using intracellular solutions, containing CsCl (133.5 mM CsCl, 1.8 mM NaCl, 1.7 mM MgCl2, 0.05 mM EGTA, 10 mM HEPES, 2 mM Mg-ATP, 0.4 mM Na2-GTP, 10 mM phosphocreatine, and 8 mM biocytin, pH: 7.2; 270–290 mOsm). To investigate isolated excitatory or inhibitory events, 5 µM SR 95531 hydrobromide (6-Imino-3-(4-methoxyphenyl)-1(6H)-pyridazinebutanoic acid hydrobromide) or 10 µM CNQX (6-Cyano-7-nitroquinoxaline-2,3-dione) and 20 µM D-AP5 (D-(−)-2-Amino-5-phosphonopentanoic acid) was added, respectively. At the end of these recordings, the identity of the recorded synaptic events was confirmed by the application of the above mentioned specific antagonists. These recordings showed no significant difference in the amplitude of excitatory events in TOR and RS cells (TOR: -43.4 ± 0.6 pA, RS: -41.6 ± 0.4 pA, *Mann-Whitney test; p = 0.1398, U = 3.42002*106, Z = 1.47654*). The simulated excitatory conductances corresponded to these events as their magnitude followed a normal distribution (mean: 0.22 nS, variance: 0.01 nS). The simulated inhibitory conductance represented both tonic and phasic inhibition (model distribution mean: 2 nS, variance: 0.1 nS) and was based on the similar inhibitory events recorded in CCK+INs (TOR: -89.1 ± 1.4 pA, RS: -78.1 ± 1.7 pA). Synaptic conductances were distributed along the dendrites and somatas of the simulated cells uniformly. In the final model, inhibitory conductances followed a uniform random temporal distribution, whereas excitatory events were aggregated into normal distributions packages at various frequencies to recreating *in vivo* relevant MP oscillations in single cells, as follows:

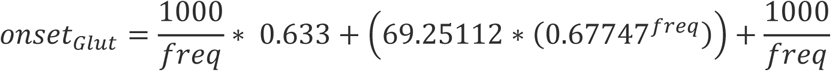

where *onset*_*Glut*_ is the timing of an individual excitatory event, and *freq* is the frequency by which excitatory packages occur. This equation is necessary for setting the width of each normal distribution according to the desired frequency, and therefore keeping the sinusoid shape of the MP during the simulated oscillations. In these simulations, 20 oscillatory cycles or in case of high frequencies at least 5 second simulation times were used. Before each run, excitatory and inhibitory event amplitudes and onsets were randomized in a unique but reproducible manner (pseudo-randomization with seed value). If the RS model (I_SA_RS) elicited APs, simulations were repeated with TOR model (I_SA_TOR+RS). The amount of excitation was set to produce firing frequencies below 50 Hz in RS cells. Changes in firing rate caused by the replacement of I_SA_RS with I_SA_TOR+RS was calculated by subtraction of the TOR firing rate from RS firing rate and normalized to the latter. Simulations were run on the Neuroscience Gateway (Sivagnanam et al., 2015).

## Data analysis and statistics

Data was analysed using Molecular Devices pClamp, OriginLab Origin, Microsoft Excel software and Python-based scripts. Normality of the data was analyzed with Shapiro-Wilks test. Data are presented as mean ± s.e.m.

## Acknowledgement

We thank Professors Henry Jerng, Paul Pfaffinger and Tõnis Timmusk for advices on DPP6 and KChIP isoforms. We are thankful for the computational resources provided by the Neuroscience Gateway. We thank Andrea Szabó, Dóra Rónaszéki, Dóra Kókay and Andrea Juszel for their their excellent technical assistance and László Barna for the kindly provided microscopy support at the Nikon Microscopy Center at the IEM HAS, which is sponsored by Nikon Europe, Nikon Austria and Auro-Science Consulting. This work was supported by Wellcome Trust International Senior Research Fellowship 087497, Hungarian Brain Research Program KTIA_13_NAP-A-I/5, European Research Council Consolidator Grant (772452, nanoAXON) to J.S., Stephen W. Kuffler Research Foundation to V.J.O., Swiss National Science Foundation (CRETP3_166815 and 31003A_170085) and from the Slack-Gyr Foundation to C.F.. ZN is the recipient of a Hungarian National Research Development and Innovation Office collaborative research grant (VKSz 14-1-2015-0155), a European Research Council Advanced Grant, and a Hungarian National Brain Research Program (NAP2.0) grant.

